# Crossover recombination and synapsis are linked by adjacent regions within the N terminus of the Zip1 synaptonemal complex protein

**DOI:** 10.1101/451674

**Authors:** Karen Voelkel-Meiman, Shun-Yun Cheng, Melanie Parziale, Savannah J. Morehouse, Arden Feil, Owen R. Davies, Arnaud de Muyt, Valérie Borde, Amy J. MacQueen

## Abstract

Accurate chromosome segregation during meiosis relies on the prior establishment of at least one crossover recombination event between homologous chromosomes, which is often associated with the meiosis-specific MutSγ complex. The recombination intermediates that give rise to MutSγ interhomolog crossovers are embedded within a hallmark meiotic prophase structure called the synaptonemal complex (SC), but the mechanisms that coordinate the processes of SC assembly (synapsis) and crossover recombination remain poorly understood. Among known central region building blocks of the budding yeast SC, the Zip1 protein is unique for its SC-independent role in promoting MutSγ crossovers. Here we report that adjacent regions within Zip1’s unstructured N terminus encompass its crossover and SC assembly functions. We previously showed that deletion of Zip1 residues 21-163 abolishes tripartite SC assembly and prevents the robust SUMOylation of the SC central element component, Ecm11, but allows excess MutSγ crossover recombination. We find the reciprocal phenotype when Zip1 residues 2-9 or 10-14 are deleted; in these mutants SC assembles and Ecm11 is hyperSUMOylated, but MutSγ crossovers are strongly diminished. Interestingly, Zip1 residues 2-9 or 2-14 are required for the normal localization of Zip3, a putative E3 SUMO ligase and pro-MutSγ crossover factor, to Zip1 polycomplex structures and to recombination initiation sites. By contrast, deletion of Zip1 residues 15-20 does not detectably prevent Zip3’s localization at Zip1 polycomplex and supports some MutSγ crossing over but prevents normal SC assembly and robust Ecm11 SUMOylation. These results highlight distinct N terminal regions that are differentially critical for Zip1’s roles in crossover recombination and SC assembly; we speculate that the adjacency of these regions enables Zip1 to serve as a liaison, facilitating crosstalk between the two processes by bringing crossover recombination and synapsis factors in close proximity to one another.

## Introduction

A unique feature of the meiotic cell cycle is how chromosomes are parsed at the first division: Homologous chromosomes (homologs) orient and precisely segregate from one another on the meiosis I spindle due to the prior establishment of recombination-based associations between homologs. Interhomolog crossover recombination creates a reciprocal splice between non-sister DNA molecules; in conjunction with sister cohesion, this DNA exchange provides a physical association between replicated homologs that is stable but nevertheless can be released to allow disjunction after the bivalent has acquired a proper orientation on the spindle [1].

Interhomolog crossovers form during meiotic prophase through the homologous recombination-based repair of a large number of programmed DSBs catalyzed by the meiosis-specific, topoisomerase-like protein, Spo11 [2]. For many organisms, the repair pathway that allows a subset of Spo11-mediated DSBs to become interhomolog crossovers involves the formation and processing of Holliday junction intermediates by meiosis-specific proteins, including the MutSγ and MutLg heterodimeric complexes which have homology to the bacterial MutS and MutL protein families, respectively [3-14]. DSB repair processes in meiotic cells also rely on meiosis-specific proteins and pathways to ensure specific desired outcomes of meiosis: for example that crossovers preferentially involve non-sister chromatids of homologous chromosomes (as opposed to involving the sister chromatids that comprise a single chromosome), and that every chromosome pair, no matter how small, receives at least one crossover [15].

Many of the meiosis-specific factors that function in MutSγ crossover recombination also serve in the assembly of a widely-conserved, prominent feature of meiotic prophase chromosomes: the synaptonemal complex (SC) [16, 17]. The SC is a proteinaceous macromolecular structure comprised largely of proteins with extensive coiled-coil (called transverse filaments) that form parallel higher order units that accumulate in an ordered fashion to create “rungs” connecting aligned chromosome axes along their entire lengths (chromosome axis structures are referred to as lateral elements in the context of the mature SC structure). As demonstrated in multiple organisms by electron and super-resolution microscopy using epitope-specific antibodies, multiple SC transverse filament protein units span the conserved 100 nm width of the SC and orient with their opposing C termini toward lateral element structures [18, 19]. In many organisms including budding yeast and mammals, a separate set of structural proteins, comprising the “central element” substructure, interface with N terminal regions of (head-to–head–oriented) transverse filament units at the midline of the SC.

While the processes of SC assembly (synapsis) and recombination are mechanistically independent and separable, nodes of crosstalk exist between the two processes. One widely-conserved example of such crosstalk is the reliance of meiotic crossover events on SC assembly proteins. Especially noteworthy is the fact that mutants missing building block components of the SC, particularly transverse filament proteins such as the budding yeast Zip1 protein, are typically deficient in MutSγ crossover formation [16]. The reliance of crossover recombination on SC proteins is perhaps unsurprising given that meiotic recombination intermediate-associated complexes fated to become chiasmata (cytological manifestations of the crossover links between homologs) embed directly within the central region of the SC [20, 21]. However we note that, at least in budding yeast, not all SC structural components play a role in crossing over and the mature SC structure itself is not a prerequisite for even MutSγ crossover recombination: Budding yeast mutants deficient in the SC central element components Ecm11 or Gmc2, or expressing an *ecm11* allele that prevents Ecm11 SUMOylation, fail to assemble tripartite SC but nevertheless exhibit *excess* MutSγ crossovers [22]. This finding indicates not only that tripartite SC is dispensable for crossing over, but that SC is associated with an activity that antagonizes interhomolog crossover formation; at least one aspect of the observed anti-crossover activity of the budding yeast SC is likely to be a capacity to inhibit Spo11 DSB formation [23, 24].

Genetic evidence from multiple systems also suggests that the recombination process directly influences SC assembly. In organisms including budding yeast and mammals, early steps in homologous recombination are a prerequisite for proper synapsis. In *spo11* mutants, which fail to initiate meiotic recombination, SC assembly does not occur extensively in mammals, or at all in budding yeast [2, 25]. In budding yeast, a subset of proteins that co-localize with MutSγ on recombining meiotic chromosomes are required downstream of DSB formation for the formation of stable, crossover-designated recombination intermediates [8, 14] and for robust SC assembly. This group of proteins includes the so–called “Synapsis Initiation Complex” factors (Zip2, Zip3, Zip4 and Spo16 [19, 26-31]. In the absence of any one of these proteins, the MutSγ heterodimer, or the SC transverse filament protein Zip1, recombination intermediates fail to form stable joint molecule (Holliday junction) structures. In the absence of Zip2, Zip4 or Spo16 (and, by default, Zip1), SC assembly is completely abolished. However, in the absence of MutSγ or the putative E3 SUMO ligase, Zip3, SC assembly is not absent but diminished, presumably due to a failure or severe delay in synapsis initiation from non-centromeric (presumed recombination-associated) chromosomal sites [26, 32]. SIC proteins have also been found to regulate the post-translational SUMOylation of SC central element component, Ecm11 [29]. Taken together, these data suggest that intermediate events in the budding yeast meiotic recombination process mediate the gradual, stepwise assembly of a recombination intermediate-associated complex that has the capacity to trigger SC elaboration. Interestingly, even in *C. elegans* where SCs assemble in the absence of recombination initiation, MutSγ-associated crossover recombination intermediates locally influence the physical and dynamic properties of the *C. elegans* SC, through a Polo-like kinase (PLK-2) signaling mechanism [33]. The observed interdependencies between synapsis and meiotic recombination indicate that the two processes are not only spatially correlated but that the mechanisms involved in each process are functionally intertwined, however we currently lack a substantial molecular understanding of the how these hallmark meiotic processes intersect.

In budding yeast it is clear that the SC transverse filament protein, Zip1, serves an early role in promoting crossover recombination independent of (and prior to) its structural role in assembling the SC; in this case, a single protein evolved dual functions to promote these unlinked but coordinated meiotic prophase processes. The budding yeast Zip1 protein thus provides an opportunity to understand how SC transverse filaments can both promote and coordinate interhomolog recombination and SC assembly. Here we present a phenotypic analysis of mutants carrying a series of non-null *zip1* alleles that encode small in-frame deletions within Zip1’s (putatively unstructured) N terminus; together with our previously-published analysis of the *zip1[Δ21-163]* mutant, these *zip1* alleles encompass distinct and nearly reciprocal phenotypes with respect to synapsis and crossover recombination. Our data identify critical N-terminal residues that correspond to Zip1’s dual function in regulating crossing over and synapsis, and suggest that these residues may encompass adjacent interaction sites for the SIC protein and putative E3-SUMO ligase, Zip3, and the SUMOylated SC central element component, Ecm11.

## Results

### MutSγ crossovers rely on residues within Zip1’s extreme N terminus

The primary amino acid sequence of Zip1 suggests that the region encompassing residues ∼175-748 of the 875 residue protein has the capacity to assemble an extended coiled-coil structure, while the flanking N-and C-terminal regions are likely unstructured. Consistent with the rod-like nature that is predicted by an extended coiled-coil, two Zip1 units assembled in a head-to-head fashion span the ∼100 nm width of the budding yeast SC central region [18, 19]; Zip1’s C termini orient toward aligned chromosome axes (lateral elements) while its N termini orient toward the central region substructure (comprised of Ecm11 and Gmc2 proteins) at the midline of the budding yeast SC. We previously reported that the non-null *zip1[Δ21-163]* mutant phenocopies SC central element-deficient *ecm11* and *gmc2* null mutants: In each of these mutants tripartite SC assembly fails but MutSγ-mediated crossover recombination events occur in excess [22]. In order to identify residues within the Zip1 and Zip1[Δ21-163] proteins that are critical for Zip1’s MutSγ-mediated crossover activity, we created and analyzed additional non-null *zip1* alleles. We found that alleles encoding disrupxtions in Zip1’s first twenty residues severely compromise Zip1’s capacity to promote MutSγ crossovers (Figure 1A).

**Fig 1.**
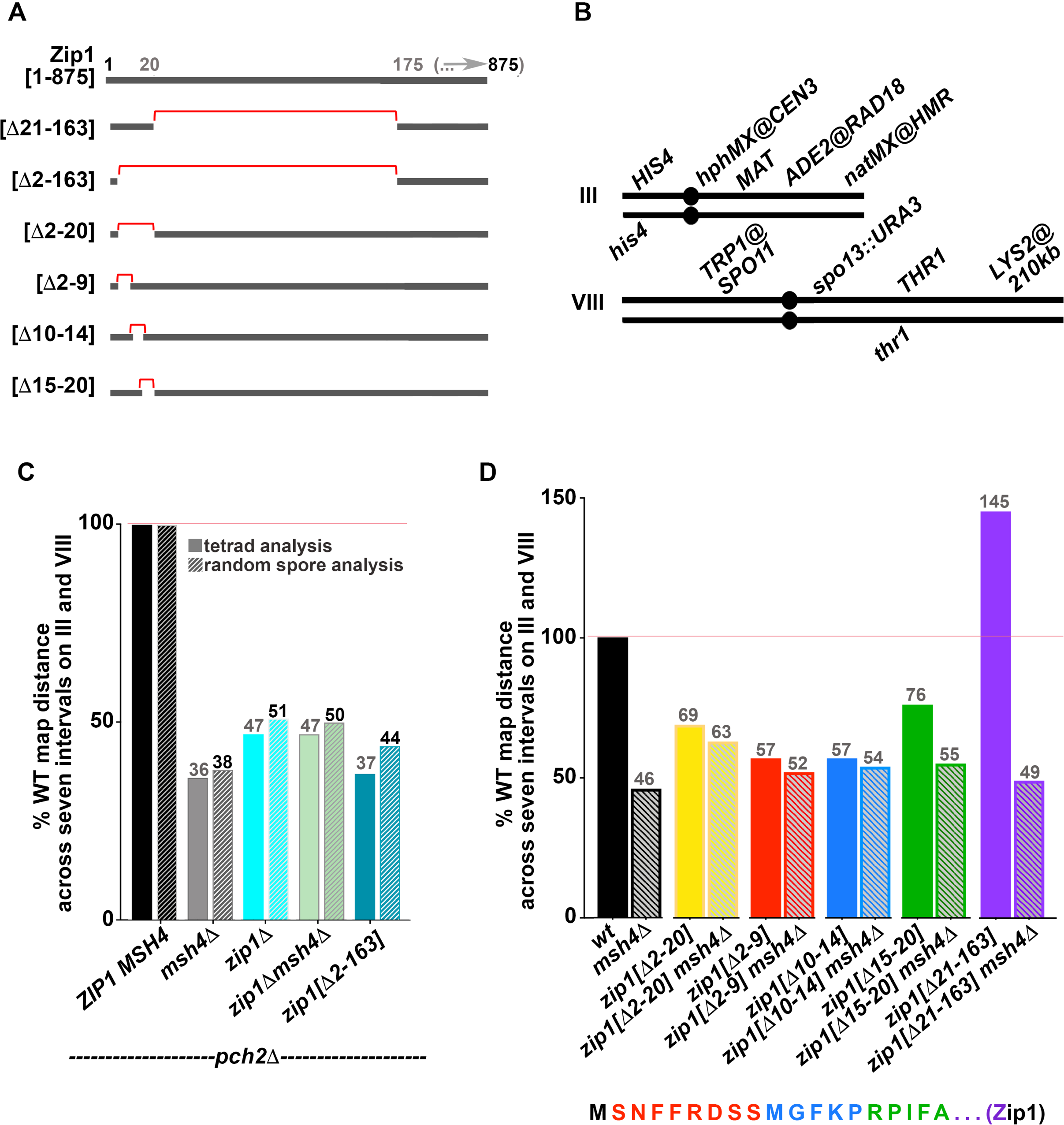
Zip1’s N terminal twenty residues are critical for MutSγ crossing over. **A)** Line illustrations represent Zip1’s N terminal region; red brackets highlight residues that are deleted in the altered versions of Zip1 analyzed in this study (precise deletion information is indicated at the left of each line).**B)** Cartoons illustrate the seven genetic intervals that were utilized to assess meiotic crossover recombination in the various strains analyzed in this study. Strains carried two distinguishable alleles of five genetic loci across the entire length of chromosome III, and of four genetic loci across more than half of chromosome VIII. Crossover recombination is measured by examining the frequency of adjacent markers becoming unlinked from one another during meiosis; a higher frequency of crossover recombination results in a larger map distance between genetic markers that define an interval. Map distances were summed across all seven intervals in each strain and these values, normalized to the wild-type value*, are plotted in **(C)** and **(D)**. Random spore analysis was used to calculate the map distances shown in the hatched bars graphed in **(C)**, while tetrad analysis was used to calculate the map distances illustrated by the solid bars in **(C, D)**; see Materials and Methods for map distance calculation procedures. Map distances for individual intervals are reported in Table 1 (*pch2* strains) and Table 2 (*PCH2* strains). Actual values corresponding to % of WT map distances (*note that “WT”= *pch2* in C) are shown in gray color above each bar in C, D. Data for wild-type, *msh4*, *zip1[Δ21-163]*, and *zip1[Δ21-163] msh4* strains were reported previously [22]. Zip1’s first twenty amino acid residues are shown below the graph in D, with colors corresponding to the residues removed in each of three zip1 mutant alleles: red = *zip1[Δ2-9],* blue = *zip1[Δ10-14]*, green = *zip1[Δ15-20].*

**Table 1.** MutSγ- and Zip1-dependent crossovers rely on Zip1’s N terminal residues. Map distances and interference values were calculated using tetrad analysis or random spore analysis and coefficient of coincidence measurements as described previously [19, 39]. 4-spore viable tetrads with no more than 2 gene conversion (non-2:2) events were included in calculations; See Table S3 for gene conversion frequencies. Table indicates map distances and their corresponding percentages of the *pch2* values for individual intervals, and the map distances and the corresponding percentage of wild type for the entire chromosome (by summing the intervals on III or VIII). For the intervals marked with (n.d.), interference measurements are not obtainable using the coefficient of coincidence method due to an absence of NPD tetrads. Data from this table is graphed in Fig. 1C.

| GENOTYPE<br>(STRAIN) | INTERVAL<br>(CHROMOSOME) | TETRAD ANALYSIS |  |  |  |  |  |  |  |  | RANDOM SPORE |  |  |  |  |  |
| --- | --- | --- | --- | --- | --- | --- | --- | --- | --- | --- | --- | --- | --- | --- | --- | --- |
|  |  | PD | TT | NPD | TOTAL | cM<br>(± SE) | % *pch2*<br>WT | cM<br>by chrm | % *pch2*<br>by chrm | NPDobs/NPDexp<br>(± SE) | # CO<br>spores | # viable<br>spores | %<br>recombinant | % *pch2*<br>WT | cM<br>by chrm | % *pch2*<br>by chrm |
| *pch2Δ*<br>(AM3724) | *HIS4-CEN3* (III) | 254 | 255 | 12 | 521 | **31.4 (2.1)** | **100** | 137.8 (III) | **100** | 0.47 (0.14) | 726 | 2714 | **26.8** | **100** | 113.4 (III) | **100** |
|  | *CEN3-MAT* (III) | 265 | 253 | 13 | 531 | **31.2 (2.1)** | **100** |  |  | 0.54 (0.16) | 713 |  | **26.3** | **100** |  |  |
|  | *MAT-RAD18* (III) | 168 | 317 | 29 | 514 | **47.8 (2.9)** | **100** |  |  | 0.55 (0.12) | 975 |  | **35.9** | **100** |  |  |
|  | *RAD18-HMR* (III) | 282 | 221 | 10 | 513 | **27.4 (2.0)** | **100** |  |  | 0.57 (0.19) | 663 |  | **24.4** | **100** |  |  |
|  | *SPO11-SPO13* (VI) | 180 | 310 | 38 | 528 | **51.0 (3.2)** | **100** | 125.6 (VIII) | **100** | 0.84 (0.16) | 988 |  | **36.4** | **100** | 92.6(VIII) | **100** |
|  | *SPO13-THR1* (VIII) | 362 | 146 | 3 | 511 | **16.1 (1.4)** | **100** |  |  | 0.46 (0.27) | 460 |  | **16.9** | **100** |  |  |
|  | *THR1-LYS2* (VIII) | 142 | 301 | 45 | 488 | **58.5 (3.6)** | **100** |  |  | 0.90 (0.17) | 1066 |  | **39.3** | **100** |  |  |
| *pch2Δ msh4Δ*<br>(AM4025) | *HIS4-CEN3* (III) | 161 | 34 | 2 | 197 | **11.7 (2.5)** | **37** | 50.7 (III) | **37** | 2.40 (1.73) | 142 | 1684 | **8.4** | **31** | 38.2 (III) | **34** |
|  | *CEN3-MAT* (III) | 179 | 22 | 1 | 202 | **6.9 (1.8)** | **22** |  |  | 3.09 (3.11) | 87 |  | **5.2** | **20** |  |  |
|  | *MAT-RAD18* (III) | 139 | 60 | 2 | 201 | **17.9 (2.6)** | **37** |  |  | 0.70 (0.50) | 235 |  | **14.0** | **39** |  |  |
|  | *RAD18-HMR* (III) | 149 | 51 | 1 | 201 | **14.2 (2.1)** | **52** |  |  | 0.51 (0.51) | 180 |  | **10.7** | **44** |  |  |
|  | *SPO11-SPO13* (VI) | 162 | 35 | 0 | 197 | **8.9 (1.4)** | **17** | 43.9 (VIII) | **35** | ND | 163 |  | **9.7** | **18** | 40.0 (VIII) | **43** |
|  | *SPO13-THR1* (VIII) | 165 | 23 | 0 | 188 | **6.1 (1.2)** | **38** |  |  | ND | 119 |  | **7.1** | **42** |  |  |
|  | *THR1-LYS2* (VIII) | 99 | 84 | 4 | 187 | **28.9 (3.4)** | **49** |  |  | 0.56 (0.29) | 392 |  | **23.3** | **59** |  |  |
| *pch2Δ zip1Δ*<br>(AM4023) | *HIS4-CEN3* (III) | 78 | 16 | 0 | 94 | **8.5 (1.9)** | **27** | 62.6 (III) | **45** | ND | 124 | 1208 | **10.3** | **38** | 46.7 (III) | **41** |
|  | *CEN3-MAT* (III) | 69 | 27 | 1 | 97 | **17.0 (3.7)** | **54** |  |  | 0.85 (0.87) | 134 |  | **11.1** | **42** |  |  |
|  | *MAT-RAD18* (III) | 62 | 28 | 2 | 92 | **21.7 (4.9)** | **45** |  |  | 1.46 (1.08) | 180 |  | **14.9** | **42** |  |  |
|  | *RAD18-HMR* (III) | 75 | 17 | 2 | 94 | **15.4 (4.8)** | **56** |  |  | 4.55 (3.33) | 126 |  | **10.4** | **43** |  |  |
|  | *SPO11-SPO13* (VI) | 56 | 40 | 1 | 97 | **23.7 (3.8)** | **46** | 61.4 (VIII) | **49** | 0.33 (0.34) | 267 |  | **22.1** | **61** | 58.9 (VIII) | **64** |
|  | *SPO13-THR1* (VIII) | 74 | 14 | 2 | 90 | **14.4 (4.9)** | **89** |  |  | 6.56 (1.80) | 122 |  | **10.1** | **60** |  |  |
|  | *THR1-LYS2* (VIII) | 53 | 36 | 1 | 90 | **23.3 (4.0)** | **40** |  |  | 0.39 (0.40) | 322 |  | **26.7** | **68** |  |  |
| *pch2Δ zip1[Δ2-163]*<br>(AM3725) | *HIS4-CEN3* (III) | 265 | 50 | 0 | 315 | **7.9 (1.0)** | **25** | 48.0 (III) | **35** | ND | 225 | 2494 | **9.0** | **34** | 44.3 (III) | **39** |
|  | *CEN3-MAT* (III) | 263 | 57 | 0 | 320 | **8.9 (1.1)** | **26** |  |  | ND | 225 |  | **9.0** | **34** |  |  |
|  | *MAT-RAD18* (III) | 203 | 104 | 5 | 312 | **21.5 (2.4)** | **45** |  |  | 0.87 (0.40) | 409 |  | **16.4** | **46** |  |  |
|  | *RAD18-HMR* (III) | 258 | 55 | 1 | 314 | **9.7 (1.4)** | **35** |  |  | 0.73 (0.73) | 246 |  | **9.9** | **41** |  |  |
|  | *SPO11-SPO13* (VI) | 226 | 82 | 2 | 310 | **15.2 (1.8)** | **30** | 50.5 (VIII) | **40** | 0.60 (0.43) | 39 |  | **13.6** | **37** | 46.0 (VIII) | **50** |
|  | *SPO13-THR1* (VIII) | 253 | 31 | 0 | 284 | **5.5 (0.9)** | **34** |  |  | ND | 201 |  | **8.1** | **48** |  |  |
|  | *THR1-LYS2* (VIII) | 155 | 121 | 8 | 284 | **29.8 (3.1)** | **51** |  |  | 0.84 (0.31) | 606 |  | **24.3** | **62** |  |  |
| *pch2Δ zip1Δ msh4Δ*<br>(AM4026) | *HIS4-CEN3* (III) | 21 | 2 | 0 | 23 | **4.3 (2.9)** | **14** | 60.2 (III) | **44** | ND | 56 | 529 | **10.6** | **40** | 46.7 (III) | **41** |
|  | *CEN3-MAT* (III) | 17 | 7 | 0 | 24 | **14.6 (4.6)** | **47** |  |  | ND | 63 |  | **11.9** | **45** |  |  |
|  | *MAT-RAD18* (III) | 14 | 9 | 1 | 24 | **31.3 (12.4)** | **65** |  |  | 1.71 (1.84) | 77 |  | **14.6** | **41** |  |  |
|  | *RAD18-HMR* (III) | 20 | 5 | 0 | 25 | **10.0 (4.0)** | **36** |  |  | ND | 51 |  | **9.6** | **39** |  |  |
|  | *SPO11-SPO13* (VI) | 11 | 10 | 0 | 21 | **23.8 (5.5)** | **47** | 62.5 (VIII) | **50** | ND | 154 |  | **29.1** | **80** | 56.0 (VIII) | **60** |
|  | *SPO13-THR1* (VIII) | 20 | 2 | 0 | 22 | **4.6 (3.1)** | **29** |  |  | ND | 47 |  | **8.9** | **53** |  |  |
|  | *THR1-LYS2* (VIII) | 12 | 9 | 1 | 22 | **34.1 (13.4)** | **58** |  |  | 1.50 (1.64) | 95 |  | **18.0** | **46** |  |  |

**Table 2.** MutSγ crossovers rely on twenty residues within Zip1’s N terminus. Map distances and interference values were calculated using tetrad analysis as described previously [39]. 4-spore viable tetrads with no more than 2 gene conversion (non-2:2) events were included in calculations; See Table S3 for gene conversion frequencies. Table indicates map distances and their corresponding percentages of wild-type values for individual intervals, and the map distances and the corresponding percentage of wild type for the entire chromosome (by summing the intervals on III or VIII). For the intervals marked with (n.d.), interference measurements are not obtainable using the coefficient of coincidence method due to an absence of NPD tetrads. Data for strains marked with an asterisk were published previously [22]. Data from this table is graphed in Fig. 1D.

| GENOTYPE<br>(STRAIN) | INTERVAL<br>(CHROMOSOME) | PD | TT | NPD | TOTAL | cM<br>(± SE) | %WT | cM<br>by chrM | %WT<br>by chrM | NPDObs/NPDexp<br>(± SE) |
| --- | --- | --- | --- | --- | --- | --- | --- | --- | --- | --- |
| WT*<br>(K842) | HIS4-CEN3 (III) | 344 | 325 | 6 | 675 | 26.7 (1.4) | 100 | 106.0 (III) | 100 | 0.19 (0.08) |
|  | CEN3-MAT (III) | 427 | 250 | 4 | 681 | 20.1 (1.2) | 100 |  |  | 0.25 (0.13) |
|  | MAT-RAD18 (III) | 255 | 405 | 14 | 674 | 36.3 (1.8) | 100 |  |  | 0.22 (0.06) |
|  | RAD18-HMR (III) | 395 | 273 | 6 | 674 | 22.9 (1.4) | 100 |  |  | 0.30 (0.13) |
|  | SPO11-SPO13 (VIII) | 251 | 401 | 21 | 673 | 39.2 (2.0) | 100 | 76.3 (VIII) | 100 | 0.35 (0.08) |
|  | SPO13-THR1 (VIII) | 565 | 94 | 1 | 660 | 7.6 (0.8) | 100 |  |  | 0.54 (0.54) |
|  | THR1-LYS2 (VIII) | 296 | 361 | 5 | 662 | 29.5 (1.3) | 100 |  |  | 0.11 (0.05) |
| msh4Δ*<br>(K852) | HIS4-CEN3 (III) | 373 | 96 | 1 | 470 | 10.9 (1.1) | 41 | 53.3 (III) | 50 | 0.35 (0.35) |
|  | CEN3-MAT (III) | 424 | 50 | 1 | 475 | 5.9 (0.9) | 29 |  |  | 1.41 (1.42) |
|  | MAT-RAD18 (III) | 275 | 183 | 7 | 465 | 24.2 (1.9) | 67 |  |  | 0.55 (0.21) |
|  | RAD18-HMR (III) | 351 | 115 | 0 | 466 | 12.3 (1.0) | 54 |  |  | n.d. |
|  | SPO11-SPO13 (VIII) | 365 | 88 | 3 | 456 | 11.6 (1.4) | 30 | 30.2 (VIII) | 40 | 1.22 (0.71) |
|  | SPO13-THR1 (VIII) | 423 | 27 | 0 | 450 | 3.0 (0.6) | 39 |  |  | n.d. |
|  | THR1-LYS2 (VIII) | 320 | 129 | 2 | 451 | 15.6 (1.4) | 53 |  |  | 0.34 (0.25) |
| zip1[Δ2-20]<br>(AM3684) | HIS4-CEN3 (III) | 463 | 107 | 3 | 573 | 10.9 (1.2) | 41 | 71.3 (III) | 67 | 1.04 (0.61) |
|  | CEN3-MAT (III) | 453 | 123 | 2 | 578 | 11.7 (1.1) | 58 |  |  | 0.52 (0.37) |
|  | MAT-RAD18 (III) | 315 | 204 | 15 | 534 | 27.5 (2.3) | 76 |  |  | 1.10 (0.30) |
|  | RAD18-HMR (III) | 346 | 197 | 6 | 549 | 21.2 (1.6) | 93 |  |  | 0.50 (0.21) |
|  | SPO11-SPO13 (VIII) | 394 | 143 | 5 | 542 | 16.0 (1.5) | 41 | 54.7 (VIII) | 72 | 0.86 (0.39) |
|  | SPO13-THR1 (VIII) | 390 | 88 | 2 | 480 | 10.4 (1.2) | 137 |  |  | 0.87 (0.62) |
|  | THR1-LYS2 (VIII) | 262 | 203 | 11 | 476 | 28.3 (2.2) | 96 |  |  | 0.69 (0.22) |
| zip1[Δ2-20] msh4<br>(K1000) | HIS4-CEN3 (III) | 365 | 58 | 3 | 426 | 8.9 (1.5) | 33 | 60.9 (III) | 57 | 2.75 (1.61) |
|  | CEN3-MAT (III) | 336 | 97 | 3 | 436 | 13.2 (1.5) | 66 |  |  | 0.94 (0.55) |
|  | MAT-RAD18 (III) | 271 | 133 | 10 | 414 | 23.3 (2.4) | 64 |  |  | 1.44 (0.47) |
|  | RAD18-HMR (III) | 297 | 117 | 2 | 416 | 15.5 (1.5) | 68 |  |  | 0.39 (0.28) |
|  | SPO11-SPO13 (VIII) | 280 | 104 | 5 | 389 | 17.2 (2.0) | 44 | 54.5 (VIII) | 71 | 1.16 (0.53) |
|  | SPO13-THR1 (VIII) | 299 | 68 | 0 | 367 | 9.3 (1.0) | 122 |  |  | ND |
|  | THR1-LYS2 (VIII) | 194 | 160 | 7 | 361 | 28.0 (2.4) | 95 |  |  | 0.52 (0.21) |
| zip1[Δ2-9]<br>(MP43) | HIS4-CEN3 (III) | 428 | 149 | 2 | 579 | 13.9 (1.2) | 52 | 72.3 (III) | 68 | 0.34 (0.24) |
|  | CEN3-MAT (III) | 441 | 140 | 1 | 582 | 12.5 (1.0) | 62 |  |  | 0.20 (0.20) |
|  | MAT-RAD18 (III) | 331 | 241 | 8 | 580 | 24.9 (1.7) | 69 |  |  | 0.44 (0.16) |
|  | RAD18-HMR (III) | 361 | 222 | 4 | 587 | 21.0 (1.4) | 92 |  |  | 0.27 (0.14) |
|  | SPO11-SPO13 (VIII) | 414 | 156 | 2 | 572 | 14.7 (1.2) | 38 | 33.7 (VIII) | 44 | 0.30 (0.22) |
|  | SPO13-THR1 (VIII) | 526 | 38 | 0 | 564 | 3.4 (0.5) | 45 |  |  | ND |
|  | THR1-LYS2 (VIII) | 400 | 165 | 2 | 567 | 15.6 (1.2) | 53 |  |  | 0.26 (0.19) |
| zip1[Δ2-9] msh4Δ<br>(MP46) | HIS4-CEN3 (III) | 389 | 106 | 4 | 499 | 13.0 (1.5) | 51 | 63.4 (III) | 60 | 1.21 (0.61) |
|  | CEN3-MAT (III) | 396 | 106 | 0 | 502 | 10.6 (0.9) | 53 |  |  | N.D. |
|  | MAT-RAD18 (III) | 299 | 185 | 9 | 493 | 24.2 (2.0) | 67 |  |  | 0.75 (0.26) |
|  | RAD18-HMR (III) | 354 | 136 | 3 | 493 | 15.6 (1.4) | 68 |  |  | 0.51 (0.30) |
|  | SPO11-SPO13 (VIII) | 364 | 117 | 3 | 484 | 14.0 (1.4) | 36 | 30.5 (VIII) | 40 | 0.70 (0.41) |
|  | SPO13-THR1 (VIII) | 452 | 24 | 0 | 476 | 2.5 (0.5) | 33 |  |  | N.D. |
|  | THR1-LYS2 (VIII) | 361 | 109 | 4 | 474 | 14.0 (1.6) | 47 |  |  | 1.70 (0.54) |
| zip1[Δ10-14]<br>(SYC107) | HIS4-CEN3 (III) | 411 | 139 | 5 | 555 | 15.2 (1.5) | 54 | 68.5 (III) | 64 | 0.94 (0.43) |
|  | CEN3-MAT (III) | 443 | 119 | 5 | 567 | 13.1 (1.4) | 75 |  |  | 1.37 (0.62) |
|  | MAT-RAD18 (III) | 336 | 219 | 5 | 560 | 22.2 (1.5) | 57 |  |  | 0.33 (0.15) |
|  | RAD18-HMR (III) | 365 | 196 | 1 | 562 | 18.0 (1.1) | 79 |  |  | n.d. |
|  | SPO11-SPO13 (VIII) | 399 | 149 | 2 | 550 | 14.6 (1.2) | 48 |  | 37 | 0.32 (0.23) |
| zip1[Δ10-14] msh4Δ<br>(SYC149) | HIS4-CEN3 (III) | 267 | 78 | 4 | 349 | 14.6 (2.0) | 52 | 61.6 (III) | 57 | 1.55 (0.79) |
|  | CEN3-MAT (III) | 286 | 65 | 4 | 355 | 12.5 (1.9) | 72 |  |  | 2.35 (1.19) |
|  | MAT-RAD18 (III) | 240 | 103 | 4 | 347 | 18.3 (2.0) | 47 |  |  | 0.82 (0.42) |
|  | RAD18-HMR (III) | 253 | 96 | 3 | 352 | 16.2 (1.8) | 71 |  |  | 0.74 (0.43) |
|  | SPO11-SPO13 (VIII) | 247 | 95 | 3 | 345 | 16.4 (1.9) | 54 |  | 42 | 0.74 (0.43) |
| zip1[Δ15-20]<br>(AF8) | HIS4-CEN3 (III) | 409 | 139 | 4 | 552 | 14.8 (1.4) | 43 | 84.3 (III) | 80 | 0.75 (0.38) |
|  | CEN3-MAT (III) | 388 | 172 | 7 | 567 | 18.9 (1.6) | 94 |  |  | 0.84 (0.32) |
|  | MAT-RAD18 (III) | 303 | 232 | 14 | 549 | 28.8 (2.1) | 79 |  |  | 0.78 (0.22) |
|  | RAD18-HMR (III) | 348 | 199 | 7 | 554 | 21.8 (1.7) | 95 |  |  | 0.58 (0.22) |
|  | SPO11-SPO13 (VIII) | 379 | 168 | 6 | 553 | 18.4 (1.6) | 47 | 54.9 (VIII) | 72 | 0.73 (0.31) |
|  | SPO13-THR1 (VIII) | 433 | 89 | 0 | 522 | 8.5 (0.8) | 112 |  |  | ND |
|  | THR1-LYS2 (VIII) | 275 | 238 | 9 | 522 | 28.0 (1.9) | 95 |  |  | 0.43 (0.15) |
| zip1[Δ15-20] msh4Δ<br>(K914) | HIS4-CEN3 (III) | 435 | 108 | 0 | 543 | 9.9 (0.9) | 35 | 63.7 (III) | 59 | n.d. |
|  | CEN3-MAT (III) | 446 | 104 | 2 | 552 | 10.5 (1.1) | 60 |  |  | 0.71 (0.50) |
|  | MAT-RAD18 (III) | 334 | 185 | 13 | 532 | 24.7 (2.2) | 63 |  |  | 1.20 (0.35) |
|  | RAD18-HMR (III) | 358 | 176 | 4 | 538 | 18.6 (1.5) | 82 |  |  | 0.42 (0.21) |
|  | SPO11-SPO13 (VIII) | 380 | 134 | 6 | 520 | 16.4 (1.7) | 54 |  | 42 | 1.14 (0.47) |
| zip1[Δ21-163]<br>(AF6) | HIS4-CEN3 (III) | 263 | 310 | 10 | 583 | 31.7 (1.8) | 119 | 139.9 (III) | 132 | 0.28 (0.09) |
|  | CEN3-MAT (III) | 241 | 329 | 13 | 583 | 34.9 (1.9) | 174 |  |  | 0.30 (0.09) |
|  | MAT-RAD18 (III) | 207 | 326 | 17 | 550 | 38.9 (2.2) | 107 |  |  | 0.35 (0.09) |
|  | RAD18-HMR (III) | 230 | 320 | 11 | 561 | 34.4 (1.9) | 150 |  |  | 0.25 (0.08) |
|  | SPO11-SPO13 (VIII) | 182 | 331 | 44 | 557 | 53.4 (3.2) | 136 | 124.7 (VIII) | 163 | 0.89 (0.16) |
|  | SPO13-THR1 (VIII) | 323 | 194 | 3 | 520 | 20.4 (1.4) | 268 |  |  |  |
|  | THR1-LYS2 (VIII) | 161 | 329 | 34 | 524 | 50.9 (3.0) | 173 |  |  |  |
| zip1[Δ21-163] msh4Δ*<br>(SYC151) | HIS4-CEN3 (III) | 481 | 116 | 2 | 599 | 10.7 (1.1) | 38 | 57.8 (III) | 54 | 0.62 (0.44) |
|  | CEN3-MAT (III) | 496 | 109 | 2 | 607 | 10.0 (1.0) | 57 |  |  | 0.73 (0.52) |
|  | MAT-RAD18 (III) | 407 | 185 | 6 | 598 | 18.5 (1.5) | 47 |  |  | 0.65 (0.27) |
|  | RAD18-HMR (III) | 397 | 199 | 4 | 600 | 18.6 (1.3) | 82 |  |  | 0.37 (0.19) |
|  | SPO11-SPO13 (VIII) | 437 | 138 | 2 | 577 | 13.0 (1.1) | 43 |  | 33 | 0.40 (0.29) |

\* Data from these strains was previously published (Voelkel-Meiman 2016)

Unlike *zip1[Δ21-163]* mutants but similar to a *zip1* null in our BR1919 (BR) strain background, *zip1[Δ2-163]* meiotic cells exhibit a severely diminished capacity to form spores (1% in *zip1[Δ2-163]* vs 5% in *zip1* vs 57% in the wild-type strain, n>1000; Table S1). The AAA ATPase protein, Pch2, is required for a meiotic checkpoint that prevents certain mutants, often those with defects in SC formation such as the *zip1* null, from progressing beyond a mid-meiotic prophase stage [34, 35]. As expected, the diminished spore formation phenotype of *zip1[Δ2-163]* is partially bypassed by removal of *PCH2* (28%; Table S1). We found that meiotic products from *pch2 zip1[Δ2-163]* strains were only slightly more viable than those from *pch2 zip1* strains (58% in *zip1[Δ2-163]* vs. 39% in *zip1*; Table S1), raising the possibility that the pro-crossover activity of Zip1 (and of Zip1[Δ21-163]) relies on Zip1 residues 2-20. Indeed, consistent with previously published data implicating Zip1 in the formation of MutSγ crossovers, genetic analysis of meiotic crossover recombination within seven intervals spanning most of chromosome III and a substantial length of chromosome VIII (Figure 1B) revealed that *pch2 msh4, pch2 zip1*, and *pch2 zip1[Δ2-163]* meiotic cells exhibit similarly diminished interhomolog crossing over relative to the *pch2* single mutant (Figure 1C, Table 1). These data in conjunction with our prior phenotypic analysis of *zip1[Δ21-163]* and *zip1[Δ21-163] msh4* [22] indicates a role for the first twenty residues of Zip1 in promoting MutSγ crossovers.

We next assessed the sporulation and interhomolog crossover recombination phenotypes of *zip1* alleles that encode proteins with smaller internal deletions of Zip1’s N terminus (Figure 1A). We found that in contrast to *zip1[Δ2-163]*, strains expressing *zip1[Δ2-9]*, *zip1[Δ10-14]*, and *zip1[Δ15-20]* sporulate as efficiently as *zip1[Δ21-163]* and wild type BR strains (57%, 54%, and 51%, respectively; Table S1). *zip1[Δ2-20]* gave an intermediate sporulation efficiency at 20%. These data, in conjunction with the *zip1* null-like sporulation efficiency of *zip1[Δ2-163]* mutants, indicate that regions 2-20 and 21-163 of Zip1 contribute redundantly to the mechanism(s) that normally prevent the *PCH2*-mediated checkpoint block to meiotic progression.

Despite their near normal sporulation efficiency and high spore viability (Table S1), *zip1[Δ2-20]*, *zip1[Δ2-9]*, *zip1[Δ10-14],* and *zip1[Δ15-20]* mutants exhibit deficiencies in MutSγ-mediated crossover recombination. Removal of the MutSγ complex protein, Msh4, reduces detectable crossovers in chromosome III and VIII genetic intervals to about 50% of wild-type levels (Figure 1D, Table 2). By contrast to *zip1[Δ21-163]* single mutants, which we previously reported exhibit elevated MutSγ-mediated crossovers [22], *zip1[Δ2-20]*, *zip1[Δ2-9]*, *zip1[Δ10-14],* and *zip1[Δ15-20]* mutants exhibit 69%, 57%, 57% and 76% of the corresponding wild-type crossover level, respectively; Figure 1D, Table 2). Importantly, removal of *MSH4* from *zip1[Δ2-20]*, *zip1[Δ2-9]*, and *zip1[Δ10-14]* strains resulted in little change in the crossover phenotype relative to their *zip1* single mutant counterparts, (*zip1[Δ2-20] msh4*, *zip1[Δ2-9] msh4*, and *zip1[Δ10-14] msh4* double mutants exhibit 63%, 52%, and 54% of wild-type crossovers, respectively) indicating that few, if any, MutSγ-mediated crossovers form in *zip1[Δ2-20]*, *zip1[Δ2-9]*, and *zip1[Δ10-14]* mutants. However, removal of *MSH4* from *zip1[Δ15-20]*, did result in a substantial further reduction in crossovers relative to the *zip1[Δ15-20]* single mutant (76% in *zip1[Δ15-20]* versus 55% in *zip1[Δ15-20] msh4*), suggesting that the zip1[Δ15-20] protein can support an intermediate level of MutSγ-mediated crossing over. The interhomolog crossover recombination phenotypes of *zip1[Δ2-9]*, *zip1[Δ10-14],* and *zip1[Δ15-20]* mutants indicate that Zip1’s first twenty residues are critical for its meiotic crossover promoting activity.

We note that each *msh4* strain missing one of these four non-null *zip1* alleles exhibits a slightly higher crossover frequency than the *msh4* single mutant. This phenomenon is particularly dramatic for the *zip1[2-20] msh4* strain, which exhibits 63% of wild type crossover frequency whereas the *msh4* single mutant exhibits 46% of wild-type crossovers (Figure 1D, Table 2). The curious result that crossovers are elevated in *msh4* mutants when Zip1’s N terminus is altered (but not when Zip1 is absent altogether; Figure 1C), raises the possibility that Zip1’s N terminal residues might normally constrain the processing of some interhomolog recombination intermediates in a manner that ensures they are MutSγ-dependent.

### Removal of residues 2-9 or 10-14 results in a novel separation-of-function Zip1 protein that is crossover-deficient but synapsis-proficient

Residues 21-163 within Zip1’s N terminal unstructured region are dispensable for MutSγ-mediated crossover recombination but essential for tripartite SC assembly [22]. We investigated whether residues 2-20 are critical for the formation of mature SC by asking whether coincident linear structures of the SC transverse filament protein (Zip1) and the SC central element protein Ecm11 assemble on surface-spread meiotic prophase nuclei from strains carrying wild-type or a mutant *zip1* allele and missing the Ndt80 transcription factor. Ndt80 is required for progression beyond a mid-meiotic prophase stage when full–length SCs are normally assembled [36], thus the *ndt80* null background allows us to maintain cells in sporulation medium for prolonged periods in order to assess the overall capacity of a strain to assemble SC. We initially examined meiotic nuclei at 24 hours after placement into sporulation medium, when ∼85% of *ZIP1 ndt80* strains in our BR genetic background exhibit nearly full synapsis [37]. We utilized a polyclonal antibody targeted against Zip1’s C terminal 264 residues [38] together with an antibody against the MYC epitope tag that is fused to the C terminus of one copy of the *ECM11* gene in these strains.

As expected based on the SC-deficient phenotype of the *zip1[Δ21-163]* mutant, meiotic prophase nuclei from the *zip1[Δ2-163]* mutant strain fail to exhibit extensive Zip1 or Ecm11-MYC linear structures on meiotic chromosomes, but instead display Zip1 or Ecm11-MYC foci of varying sizes, sometimes accompanied by a large “polycomplex” aggregate of these SC central region proteins (Figure S1). Interestingly, *ndt80* meiotic cells expressing *zip1[Δ2-20]* also fail to exhibit any detectable SC formation, even after 24 hours in sporulation medium. These data in conjunction with the deficient SC assembly phenotype of *zip1[Δ21-163]* [22] indicate that residues within both the 2-20 and 21-163 regions of Zip1 are required for Zip1’s capacity to assemble SC.

However, we found that not all residues within Zip1’s 2-20 region are critical for SC assembly. Surface–spread meiotic prophase nuclei in *zip1[Δ2-9]* and *zip1[Δ10-14]* strains often display extensive SC, as reflected by long linear assemblies of coincident anti-Zip1 and anti-Ecm11-MYC label (Figure 2). We also observed meiotic nuclei with extensive SCs in a strain expressing *zip1[10-14→A]*, where Zip1’s residues 10-14 are replaced with alanine (Figure 2).

**Fig 2.**
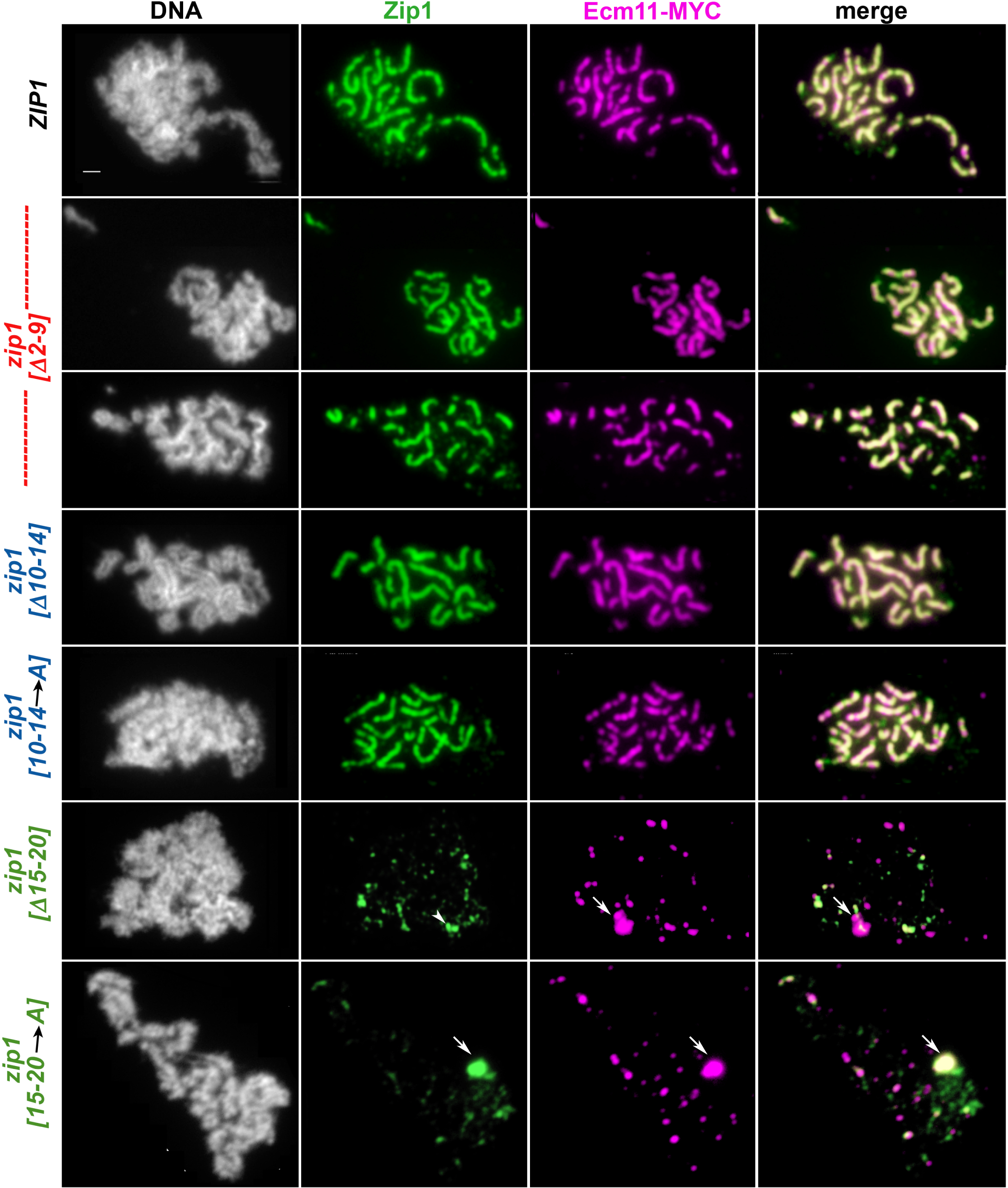
SC assembly requires Zip1 residues 15-20 but not residues 2-14. Panels show representative surface-spread mid-meiotic prophase nuclei from wild-type (top row), and various *zip1* internal deletion or residue-substitution alleles (genotypes indicated at left). Note that strains carrying a short internal deletion of Zip1 residues 10-14 or 15-20 exhibit a similar phenotype, respectively, as strains in which Zip1 residues 10-14 or 15-20 are replaced with alanine. All strains carry an *ndt80* null allele, which allows meiotic cultures to accumulate at mid-late prophase stages when full-length SCs are normally present. Mid-meiotic prophase chromosomes are stained with DAPI to label DNA (white), anti-Zip1 to label Zip1 (green), and anti-MYC to label Ecm11 (magenta). The merge between Zip1 and Ecm11 channels is shown in the final column. Quantitation of the number and cumulative length of SC linear assemblies in wild type as well as each internal deletion *zip1* mutant strain is given in Fig. 3. Arrows point to polycomplex structures. Arrowhead indicates a large focus or pair of foci that measures at 0.7 µm and thus may have been included in the assessment of “linear” SC structures (see Fig. 3). Scale bar, 1 μm.

The tripartite nature of the mature SC in budding yeast is reflected by the orientation of Zip1 transverse filament proteins relative to the central element substructure, whereby Zip1 C termini orient toward homologous axes and away from the central element, which is positioned at the midline of the SC. This tripartite organization can be detected using structured illumination microscopy (SIM) on surface-spread meiotic chromosomes labeled with antibodies targeting the C terminal region of Zip1 and SC central element components [19]. With the increased resolution that SIM affords, our anti-Zip1 antibody localizes as a wide ribbon on linear SC structures; one can often observe parallel tracts of Zip1 C termini flanking the central element protein(s) within subsections of such a Zip1 linear element. We used SIM on surface-spread meiotic chromosomes to ask whether the SC structures in *zip1[Δ2-9]* meiotic nuclei have this canonical, tripartite organization. We observed no detectable difference in the organization of Zip1 and the central element proteins within SCs assembled by wild type Zip1 versus Zip1[Δ2-9] protein: Antibodies targeting the C terminus of Zip1 were observed to flank the SC central element substructure (labeled with antibodies directed against the Ecm11 and/or Gmc2 proteins; see Methods) within SCs assembled by Zip1[Δ2-9] (Figure S2).

These observations indicate that, in reciprocal fashion to residues 21-163, residues 2-15 are required for Zip1’s MutSγ crossover-promoting activity but are dispensable for its capacity to assemble tripartite SC.

In contrast to the robust synapsis observed in *zip1[Δ2-9]* and *zip1[Δ10-14]* strains, extensive coincident linear assemblies of Zip1 and Ecm11 were not detectable in meiotic prophase nuclei from *zip1[Δ15-20]* or *zip1[15-20* à*A]* mutant strains (Figure 2). Instead, the vast majority of meiotic prophase nuclei from these strains exhibit foci of Zip1 and Ecm11 of varying sizes, which are sometimes, but not always, coincident.

Our phenotypic assessment of three novel non-null *zip1* alleles thus indicates that Zip1’s first twenty residues correspond to adjacent regions that function somewhat independently of one another: residues 2-14 are essential for normal MutSγ crossovers but dispensable for SC assembly, whereas residues 15-20 (and certainly residues 21-163; [22]) are less critical for MutSγ crossovers but crucial for SC assembly.

### SC assembles early MutSγ crossover-deficient but SC-proficient *zip* mutants

To gain a deeper understanding of the extent of SC assembly in *zip1[Δ2-9]*,*zip1[Δ0-14]*, and *zip1[Δ15-20]*,strains, we examined 50 meiotic surface spread nuclei from each strain at 15, 18, 21 and 24 hours after placement into sporulation medium. In addition to a wild-type control strain, our analysis also included the *zip3* mutant, which resembles *zip1[Δ2-9]* and *zip1[Δ10-14]* mutants in its failure to form MutSγ crossovers but proficiency for SC assembly (albeit diminished). We quantified the extent of SC assembly in each of these strains at each time point by measuring the number of linear assemblies of Zip1, Ecm11-MYC, or coincident Zip1-Ecm11-MYC (Figure 3, left column) and the cumulative length of Zip1, Ecm11-MYC, and coincident Zip1-Ecm11-MYC linear structures (Figure 3, center column).

**fig 3.**
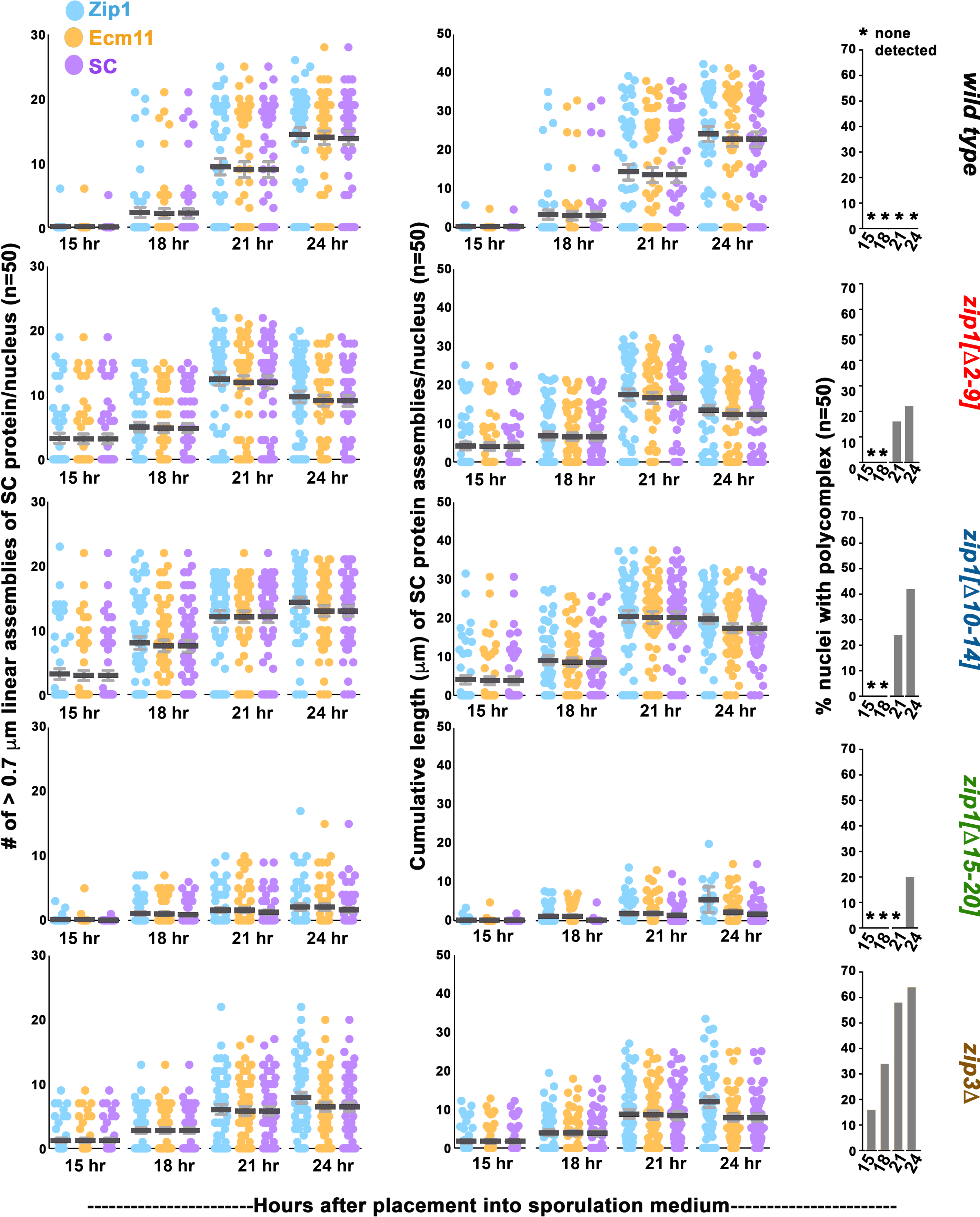
SC assembly requires Zip1 residues 15-20 but not residues 2-14. Each dot in the scatterplots at far left represents the number of linear assemblies of Zip1 (blue). Ecm11 (orange) or coincident Ecm11 and Zip1 (purple) detected in surface spread meiotic nuclei of wild-type, *zip1* or *zip3* mutant strains (genotype indicated at the far right), at 15, 18, 21 or 24 hours after placement into sporulation medium (time points indicated on the *x* axis). 50 nuclei were examined for each strain at every individual time point. Assemblies of SC proteins were considered to be linear if they measured 0.7 µm or greater in length, although some large or adjacent foci potentially were included in these calculations (see Fig. 2 legend). Individual dots in the middle column of scatterplots indicate the cumulative length of the linear assemblies of Zip1 (blue), Ecm11 (orange) or coincident Ecm11 and Zip1 (purple, “SC”) detected in the surface spread nuclei of indicated strains. Dark and light grey bars indicate the mean, and standard error of the mean, respectively. Bar graphs at far right indicate the percentage of nuclei in these datasets that exhibited a Zip1 polycomplex aggregate (see Fig. 2). The vast majority of polycomplex structures detected contained both Zip1 and Ecm11 proteins. An asterisk indicates that zero polycomplex structures were detected in any nuclei at the indicated time point.

Our analysis revealed that SC cumulative length per meiotic nucleus (as measured by coincident Zip1 and Ecm11-MYC; purple dots in Figure 3’s center scatterplots) was only slightly reduced in *zip1[Δ2-9]* and *zip1[Δ10-14]* meiocytes relative to wild type: *zip1[Δ2-9]* and *zip1[Δ10-14]* populations of meiotic nuclei exhibited a maximum of 32 and 38 microns, and a on average 17 and 20 microns, of cumulative SC length per nucleus, respectively, at the time point displaying the most abundant SC, whereas *ZIP1* meiotic nuclei exhibited a maximum of 41 and average of 23 microns at the time point with the most abundant SC. *zip3* mutants exhibited a more dramatic reduction in SC cumulative length relative to wild type, with a maximum of 25 and a mean of 8 microns at the time point exhibiting most abundant SC (Figure 3, center column).

Interestingly, SC cumulative length was highest for wild-type populations at the 24 hour time point, while SC cumulative length in *zip1[Δ2-9], zip1[Δ10-14]*, and *zip3* strains peaked at the 21 hour time point. Moreover, at the earliest time point (15 hour) both the number of SC linear assemblies and SC cumulative length per meiotic nucleus was significantly higher in *zip1[Δ2-9], zip1[Δ10-14]* and *zip3* populations (exhibiting a maximum of 25, 31, 12 microns and mean of 4, 4, 2 microns, respectively) relative to the *ZIP1 ZIP3* control population, which exhibited a mean of 0.1 micron of cumulative SC length per nucleus). This difference in SC accumulation at the earliest time point in this experiment suggests that SC initiates earlier and/or assembles faster than wild type in *zip1[Δ2-9], zip1[Δ10-14]*, and *zip3* meioses.

SC assembly appeared less extensive at the 24 hour time point in the *zip1[Δ2-9]*, and *zip1[Δ10-14]* mutants (Figure 3). Although the relative decrease in cumulative SC at 24 hours compared to the 21 hour time point is not dramatic, both the range and average cumulative SC length per nucleus shifted lower at the 24 hour time point in *zip1[Δ2-9]*, and *zip1[Δ10-14]* strains. Furthermore, Zip1 polycomplex aggregates, while undetectable in 100 *zip1[Δ2-9]* and *zip1[Δ10-14]* meiotic nuclei from 15 and 18 hour time points, were observed at the 21 hour time point and were observed even more frequently (30-40% of nuclei, n=50) at the 24 hour time point (Figure 3, right column). As polycomplex structures tend to form in situations when SC assembly is disrupted, the sub-peak level of assembled SC at the 24 hour time point and the increased occurrence of polycomplex structures together suggest the possibility that SCs assembled in *zip1[Δ2-9] ndt80* and *zip1[Δ10-14] ndt80* strains are less stable than SCs assembled in *ZIP1 ndt80* strains. The early appearance of polycomplex structures and overall lower extent of assembled SC observed in the *zip3* mutant (ranging from 1.7 microns at 15 hours to 8.4 microns at the time point with peak SC assembly, 21 hours) also raises the possibility that SCs assemble early but may be unstable when Zip3 is absent.

Our time course experiment additionally confirmed that *zip1[Δ15-20]* meiotic nuclei fail to assemble extensive SC at any point during meiosis progression (prior to the *ndt80* late meiotic prophase arrest). Out of the 200 *zip1[Δ15-20]* meiotic nuclei analyzed over the time course, zero exhibited robust long linear structures containing coincident Zip1 and Ecm11-MYC. However, large or adjacent foci of coincident Zip1 and Ecm11 were sometimes included in our SC measurements (which recorded any Zip1 or Ecm11 continuous structures with a dimension of 0.7 micron or more), and occasionally a meiotic nucleus displayed linear elements of Ecm11 and Zip1 with a frayed and diffuse appearance (Figure S3). The average cumulative length of SC per nucleus detected in *zip1[Δ15-20]* populations was 1.7 microns at the peak time point (24 hours; Figure 3). Zip1 polycomplex structures were observed only at the 24 hour time point in *zip1[Δ15-20]* strains (Figure 3, right column).

### Crossover-deficient, synapsis-proficient *zip1* mutants initiate SC assembly from both centromeric and non-centromeric chromosomal sites

While the earliest SC assembly events that occur during meiosis in budding yeast have been found to preferentially initiate at centromeres [32], SC assembly events are also associated with non-centromeric sites (presumably recombination sites) in wild-type meiotic nuclei at intermediate stages of synapsis. In the *zip3* mutant, by contrast, new SC assembly events associate predominantly with centromeres at both early and later meiotic prophase stages [32]. As the SC assembly and meiotic crossover phenotypes in *zip1[Δ2-9]* and *zip1[Δ10-14]* strains resemble the phenotypes found in *zip3*, we asked whether new SC assembly events in *zip1[Δ2-9]* and *zip1[Δ10-14]* meiotic nuclei associate with centromeres more often than wild-type nuclei at intermediate stages of synapsis. Surface-spread meiotic chromosomes from 15, and 18 hour time points were co-labeled with antibodies that target Zip1, and that target the MYC epitope that is fused to the Ctf19 centromere protein in these strains. In order to enrich for new, singular SC assembly events in our analysis, we identified the total number of Zip1 linear stretches measuring between 0.7-1.0 micron in length, and asked which are directly adjacent to (overlapping) a Ctf19-MYC focus. We found that 83% (29 out of 35) of such short SC structures were associated with a Ctf19-MYC focus in *zip3* meiotic nuclei, consistent with previously–published data. By contrast, 48%, 52% and 64% (35/73, 27/52, and 23/36) of short Zip1 linear structures were associated with Ctf19-MYC *zip1[Δ2-9], zip1[Δ10-14]*, or *ZIP1 ZIP3* meiocytes, respectively (Figure S4). Thus, in contrast to *zip3* mutants, *zip1[Δ2-9],* and *zip1[Δ10-14]* mutants display a robust capacity to initiate SC assembly from non-centromeric sites on meiotic chromosomes.

### Adjacent regions within Zip1’s first twenty residues have opposing effects on the SUMOylation of an SC central element protein

Humphryes et. al (2013) demonstrated that SUMOylated Ecm11 is required for SC assembly and that Zip1 along with SIC proteins Zip2, and Zip4 (but not Zip3) are required for a threshold level of Ecm11 SUMOylation during meiosis [29]. This report also revealed that Ecm11 is hyper-SUMOylated in mutants missing the putative SUMO E3 ligase and SIC protein, Zip3. To ask whether the N terminal twenty residues of Zip1 regulate Ecm11 SUMOylation, we evaluated the abundance of Ecm11 forms in meiotic extracts from *ndt80* strains homozygous for *ZIP1*, *zip1[Δ2-9], zip1[Δ10-14]*, or *zip1[Δ15-20]*, a *zip1* null, or a *zip3* null allele.

A Western blot can readily detect three forms of Ecm11-MYC in protein extracts from meiotic cells homozygous for MYC-tagged Ecm11 [19, 29, 39]. UnSUMOylated Ecm11-MYC migrates near the 75 kD marker on a protein gel, whereas monoSUMOylated and polySUMOylated Ecm11-MYC is positioned near the 100 kD and 150 kD positions, respectively. HyperSUMOylated Ecm11-MYC, which is found in *zip3* meiotic extracts, migrates at various positions between the 150 kD and 250 kD markers (Figure 4A).

**fig 4.**
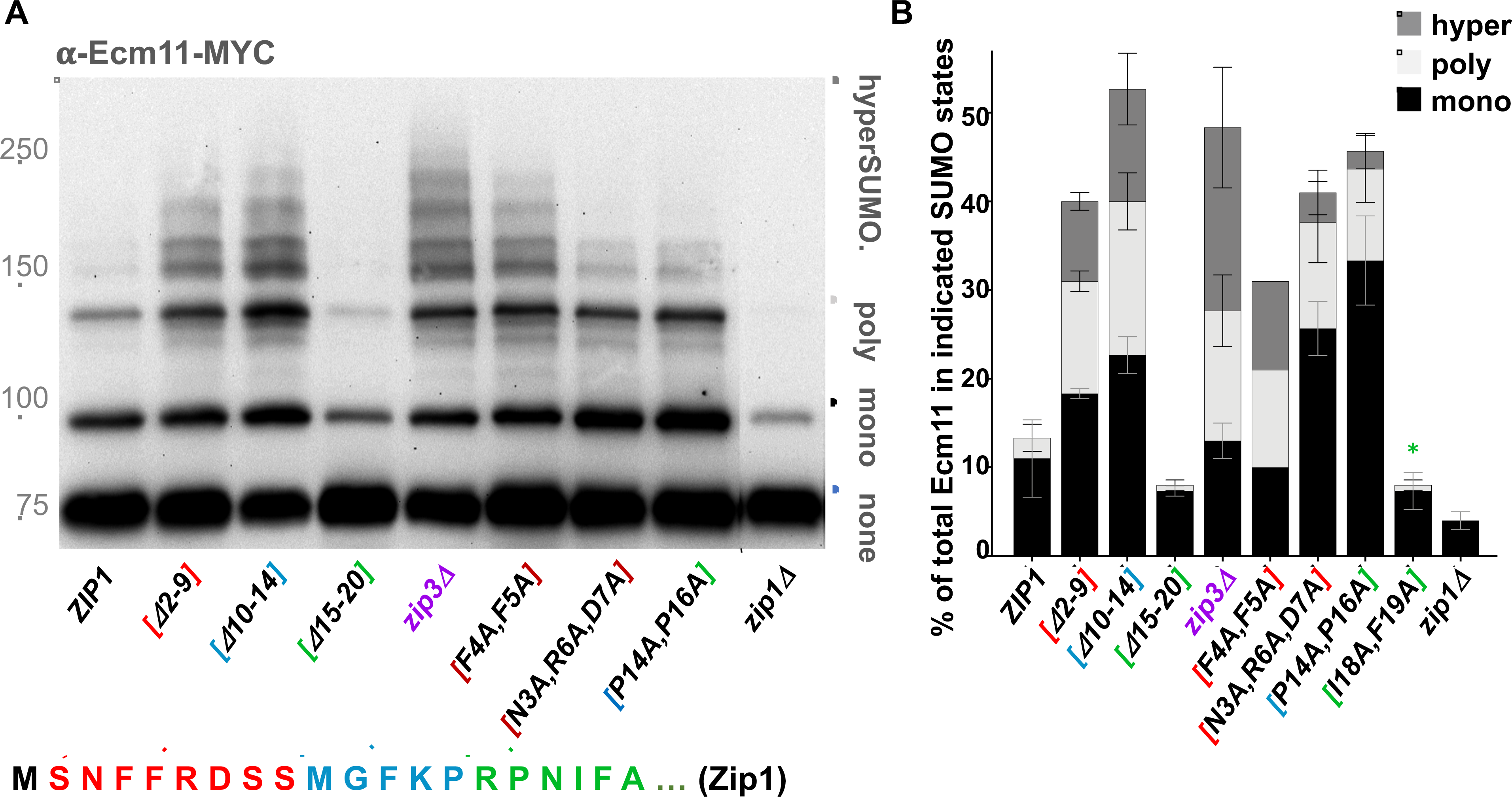
Zip1 residues 15-20 promote, while residues 2-14 limit, SUMOylation of the SC central element protein Ecm11. A) A representative Western blot using an anti-MYC antibody reveals unSUMOylated, mono-SUMOylated, poly-SUMOylated and hyper-SUMOylated forms of Ecm11-MYC in meiotic extracts prepared from *ZIP1 ZIP3*, *zip3*, or various *zip1* mutant strains (protein alterations caused by each *zip1* allele is indicated on the *x* axis). All strains carry an *ndt80* null allele, which causes a meiotic arrest that ensures maximal enrichment of midmeiotic prophase stage cells at 24 hours after placement into sporulation medium [37]. Meiotic extracts were prepared as previously described at the 24 hour time point [19, 29]. **B)** Graph plots the percentage of mono-SUMOylated (dark shaded bar), poly-SUMOylated (light bar), or hyper-SUMOylated (grey shaded bar) forms of Ecm11-MYC detected in each strain at the 24 hour time point. The absence of a value for any strain indicates that this form of Ecm11 was detected in less than 1% of the total population. Error bars represent the range of values from three independent meiotic cultures, except for in the case of the *zip1[F4A, F5A]* strain, where only one experiment was performed. * indicates a strain not shown in the blot in (A).

We found the proportion of SUMOylated Ecm11-MYC in *ZIP1 ZIP3 ndt80* meiotic extracts at the 24 hour time point to be, on average, 14%, wherein 11% of total Ecm11-MYC was in the monoSUMOylated form and 3% was in the polySUMOylated form (three replicates; Figure 4B). Consistent with prior results [19, 29], *zip1* null strains exhibited a relatively low level of SUMOylated Ecm11: An average of 4% of total Ecm11-MYC was in the monoSUMOylated form, while polySUMOylated Ecm11-MYC was below levels of detection (less than 1%) 24 hours after placement into sporulation medium (Figure 4B). Again consistent with prior findings [29], *zip3* meiotic extracts exhibited not only an elevated level of polySUMOylated Ecm11-MYC (15% of total Ecm11-MYC, on average, over three replicates; Figure 4B), but also exhibited an abundance of hyperSUMOylated Ecm11 (19% of total Ecm11-MYC, on average, over three replicates; Figure 4B).

We found that residues 15-20 are important for Zip1’s capacity to promote Ecm11 SUMOylation. In *zip1[Δ15-20]* meiotic extracts at the 24 hour time point, on average only 8% of total Ecm11 was in a SUMOylated form, wherein 7% of total SUMOylated Ecm11-MYC detected was monoSUMOylated and 1% polySUMOylated (Figure 4B). Given the SC assembly defect of *zip1[Δ15-20]* strains, this result bolsters a direct correlation between Ecm11 SUMOylation and SC assembly.

In striking contrast, cells expressing the *zip1[Δ2-9]* or *zip1[Δ10-14]* alleles exhibit robust levels of Ecm11 SUMOylation as well as the hyperSUMOylated forms of Ecm11 that are characteristic of the *zip3* mutant. At the 24 hour time point, an average of 40% of Ecm11-MYC was SUMOylated in *zip1[Δ2-9]* meiotic extracts, with 18% in the monoSUMOylated form, 13% in the polySUMOylated form, and 9% in the hyperSUMOlated form (Figure 4B). Likewise, an average of 54% of Ecm11-MYC was SUMOylated in *zip1[Δ10-14]* meiotic extracts, with 23% in the monoSUMOylated form, 18% in the polySUMOylated form, and 13% in the hyperSUMOylated form (Figure 4B).

It has been proposed that the hyperSUMOylated forms of Ecm11 that occur in *zip3* mutant meiotic cells correspond to Ecm11 linked to poly-SUMO branched chain structures of various sizes and shapes [29, 40]. The accumulation of such extensively SUMOylated Ecm11 protein in *zip1[Δ2-9]* and *zip1[Δ10-14]* mutants indicates that residues within the 2-14 region of Zip1 are dispensable for Ecm11 SUMOylation *per se*, but regulate the extent and/or manner of Ecm11 SUMOylation. Zip1’s residues 2-14 appear to control similar aspects of Ecm11 SUMOylation as the putative SUMO E3 ligase, Zip3, raising the possibility that Zip1’s N terminal residues regulate Ecm11 SUMOylation in part through an interaction with Zip3.

### The phenotype of two-or three-residue alterations within Zip1’s N terminus resemble corresponding small internal deletion alleles

We additionally found that *zip1* alleles encoding proteins with alanine substitutions in place of dual or triple residues within Zip1’s N terminus exhibit the distinguishing Ecm11 SUMOylation and synapsis phenotypes of corresponding internal deletion *zip1* alleles. *zip1[N3A, R6A, D7A]*, *zip1[F4A, F5A]*, and *zip1[P14A, P16A]* exhibited elevated levels of SUMOylated Ecm11 during meiosis, reminiscent of the *zip1[Δ2-9]* and *zip1[Δ10-14]* mutants, although we note that *zip1[N3A, R6A, D7A]* and *zip1[P14A, P16A]* strains exhibit a particular abundance of monoSUMOylated relative to polySUMOylated and hyperSUMOylated Ecm11, which differs slightly from the distribution of SUMOylated Ecm11 forms in *zip1[F4A, F5A], zip1[Δ2-9], zip1[Δ10-14]* or *zip3* mutants (Figure 4B). By contrast, meiotic extracts from *zip1[I18A, F19A]* strains exhibit a dramatic reduction in SUMOylated Ecm11, reminiscent of meiotic extracts from *zip1[Δ15-20]* and *zip1* null strains (Figure 4B).

We furthermore found that *zip1[N3A, R6A, D7A]* and *zip1[I18A, F19A]* exhibit SC assembly phenotypes that are generally reminiscent to corresponding internal deletion *zip1* alleles. Linear stretches of coincident Zip1 and Ecm11-MYC were often detectable on surface-spread meiotic chromosomes from *zip1[N3A, R6A, D7A] ndt80* strains at multiple time points in a meiotic time course, while such extensive SC structures were absent from meiotic chromosomes in *zip1[I18A, F19A] ndt80* strains at all time points. Interestingly, *zip1[F4A, F5A]* strains exhibit the capacity to assemble SC, but to a lower extent than *zip1[Δ2-9]* or *zip1[N3A, R6A, D7A]* strains (Figure S5), suggesting that the Zip1[F4A, F5A] protein has a negative activity that renders it less capable (relative to Zip1[Δ2-9] or *zip1[N3A, R6A, D7A]*) of supporting SC assembly.

We also found that meiotic crossovers are reduced in *zip1[N3A, R6A, D7A], zip1[F4A, F5A],* and *zip1[I18A, F19A]* point mutants, to a level that is intermediate between wild-type and the corresponding deletion strains (Table S2). Interestingly, *zip1[F4A, F5A]* exhibited the most dramatic deficit in meiotic crossovers, particularly on chromosome VIII.

The phenotypes of these *zip1* point mutants support the idea that while Zip1’s first twenty residues encompass both crossover recombination and SC assembly functionalities, adjacent sites within this region maintain different and independent roles in regulating synapsis.

### Residues within the 2-14 region influence Zip1’s capacity to interface with Zip3 at polycomplex structures

The shared phenotypes of *zip1[Δ2-9], zip1[Δ10-14]* and *zip3* mutants made us wonder whether the N terminus of Zip1 directly or indirectly interacts with the Zip3 protein. Prior evidence for an interaction between Zip1 and Zip3 includes the observation that Zip3 is detected throughout Zip1 polycomplex structures that assemble in contexts where SC assembly fails [26, 32]. To explore the possibility that Zip1’s N terminus mediates an interaction with the Zip3 protein, we examined the distribution of Zip3 at Zip1 polycomplex structures assembled in *spo11* meiotic cells, which fail to initiate recombination and thus also SC assembly [41, 42].

Of the polycomplex structures assembled by wild-type Zip1 and zip1[Δ15-20], 100% (50/50) exhibited Zip3-MYC distributed uniformly across the entire structure (Figure 5). Frequently, additional “capping” structures of coincident Zip3-MYC and Zip4-HA protein flank the Zip1 polycomplex in these *spo11* meiotic nuclei. Intriguingly, however, among 50 meiotic nuclei examined from *spo11 zip1[Δ2-9]* and *spo11 zip1[Δ10-14]* strains, Zip3-MYC was completely absent from the bulk of the Zip1 polycomplex structure (Figure 5). Instead, Zip3-MYC co–localized with Zip4-HA in the capping configuration at opposite ends of the polycomplex. Often these “capping” structures of coincident Zip3-MYC and Zip4-HA were observed at a substantial distance away from the polycomplex aggregate of Zip1 protein.

**fig 5.**
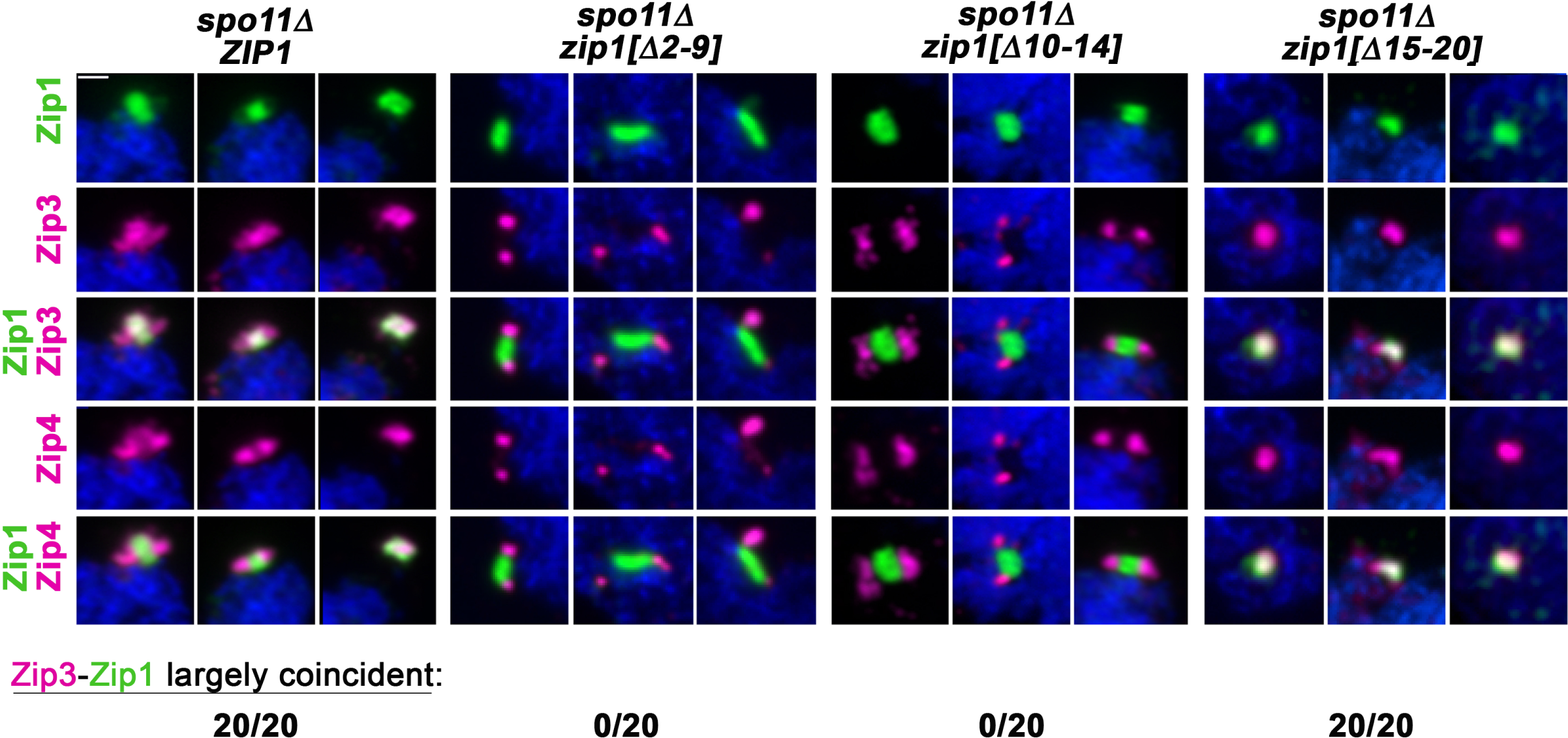
Zip1 residues 2-14 promote the localization of Zip3 to Zip1 polycomplex structures. Four groups of panels correspond to three representative images of Zip1 polycomplex structures (first row, green) in *spo11* meiotic prophase nuclei expressing either wild-type *ZIP1* (far left group), *zip1[Δ2-9]* (second group), *zip1[Δ10-14]* (third group), or *zip1[Δ15-20]* (far right group). The localization of Zip3-MYC (second row, magenta) and Zip4-HA (fourth row, magenta) is also assessed on these surface-spread meiotic prophase nuclei. Merged images between either Zip3-MYC and Zip1 or Zip4-HA and Zip1 are shown in the third and fifth rows, respectively. Scale bar, 1 µm. The number of polycomplexes (n=20) for which Zip3 is nearly fully coincident with Zip1 is displayed at the bottom of the corresponding strain’s images (n=20).

We furthermore found that Zip3 is diminished at polycomplexes assembled by Zip1[N3A, R6A, D7A] or Zip1[F4A, F5A] protein (Figure S5), although the absence of Zip3 from the bulk of these Zip1 polycomplex structures is less dramatic than what is observed at Zip1[Δ2-9] or zip1[Δ10-14] polycomplex. Finally, as expected based on the robust localization of Zip3 to zip1[Δ15-20] polycomplex, we found that Zip3 also localizes uniformly throughout polycomplexes built of Zip1[Δ21-163] protein (Figure S5).

These data indicate that, at least in the context of polycomplex structure, Zip1’s residues 2-14 mediate a direct or indirect interaction with Zip3.

### Residues within Zip1’s 2-14 region are critical for Zip3 recruitment to recombination initiation sites

Zip3 and other SIC proteins (such as Zip2, Zip4 and Spo16) form foci that co-localize with MutSγ along aligned homologous chromosomes at mid meiotic prophase [26]. Consistent with the notion that such Zip3 foci mark recombination intermediates, Zip3 has been detected at multiple DSB hotspots using Chromatin Immunoprecipitation (ChIP) in conjunction with quantitative PCR (qPCR) [43]. Zip1 was found to be required for the recruitment of Zip3 to the DSB sites examined, thus we asked whether the capacity of Zip1 to recruit Zip3 to DSB sites relies on the N terminal residues of Zip1 that facilitate Zip3’s localization to Zip1 polycomplex.

We performed ChIP and qPCR on meiotic cell extracts from *ZIP1*, *zip1[Δ2-9]*, *zip1[Δ10-14]*, and *zip1* null strains expressing a Zip3 protein with three copies of the FLAG epitope fused to its C terminus. Strains for this experiment were built in the SK1 genetic background, to ensure maximal synchrony over a meiotic time course (SK1 strains enter and/or progress through meiosis more synchronously than the BR strain background). ChIP-qPCR was performed at multiple time points during sporulation, and the time course experiment was performed in triplicate for each strain except for the *zip1* null negative control, where the single experiment performed gave results that are consistent with prior published data [43].

We examined Zip3-6xHIS-3xFLAG association with chromosomal sites corresponding to three known DSB hotspots, centromeres, or the chromosome axis [43]. Sequences enriched for Rec8 that are embedded in the proteinaceous chromosome axis are generally anti-correlated with DSB sites, but may associate with DSB repair intermediates, according to a “loop-tether” model for DSB formation in budding yeast [44]. A sequence internal to the large *NFT1* open reading frame was previously found to be devoid of Zip3 binding [45, 46] and thus served as a normalization control for Zip3 relative enrichment values.

In strains carrying wild-type *ZIP1,* Zip3 was detected in higher abundance at meiotic centromeres relative to axis or DSB sites within two hours after placement of into sporulation medium, consistent with previously published results. Between two to four hours after placement into sporulation medium, Zip3 became enriched at a chromosome axis site as well as at three DSB hotspots (*GAT1*, *BUΔ23*, and *ERG1*; Figure 6). Zip3 localization to all sites peaked at the 4 hour time point in *ZIP1* meiotic cells, which corresponds to maximal DSB activity at the *BUΔ23* locus in this SK1 strain background [43]. At this four hour time point, *GAT1* and *BUΔ23* DSB sites exhibited a more than 20-fold enrichment for Zip3 relative to the *NFT1* control (Figure 6). Consistent with DSB repair timing in this genetic background, Zip3 enrichment at all sites dramatically diminished between four to six hours and was at pre-meiotic levels by eight hours after placement in sporulation medium. Consistent with prior findings, Zip3 was virtually undetectable at DSB, axis and centromere sites in the *zip1* null strain (Figure 6).

**fig 6.**
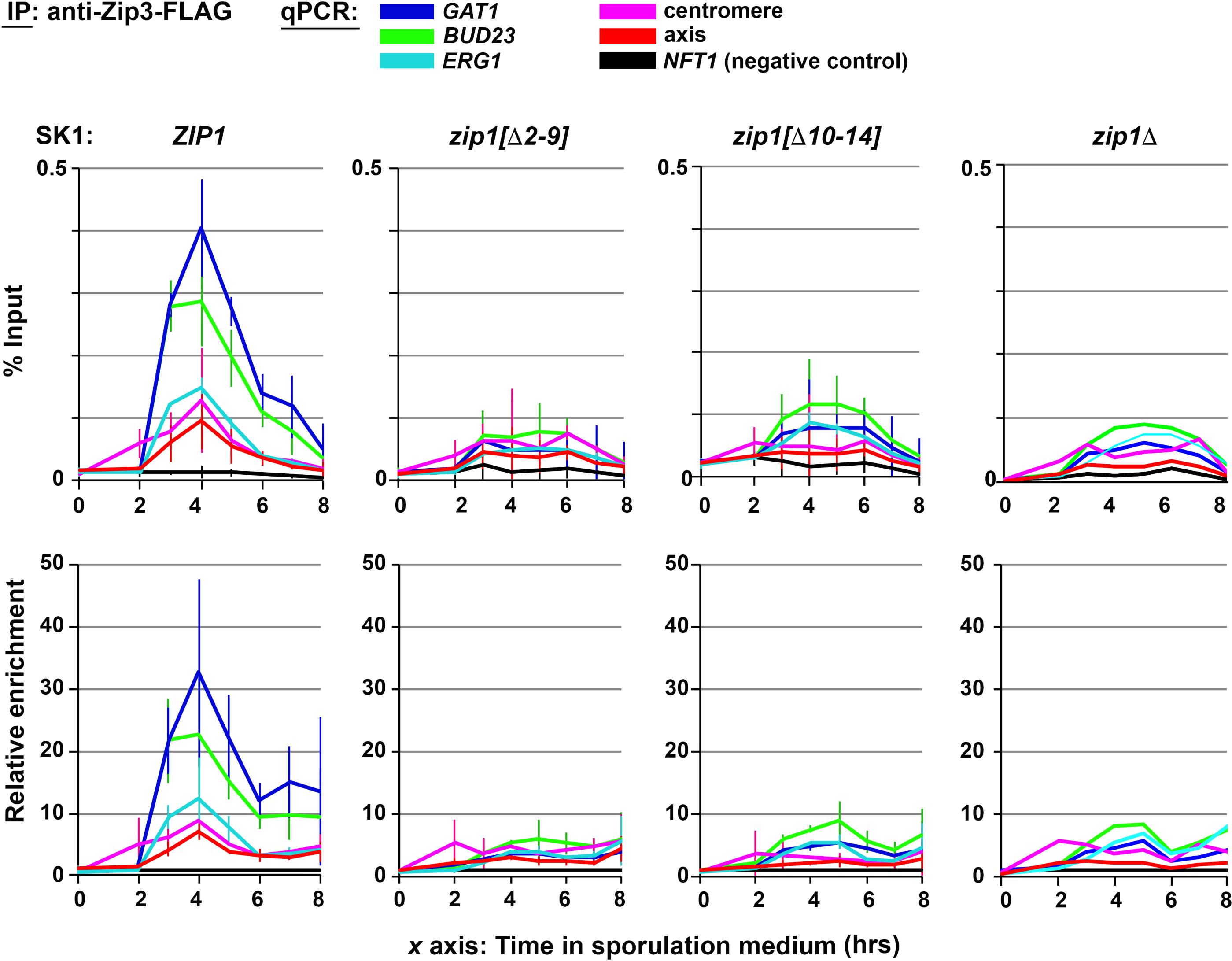
Residues 2-14 are required for Zip1’s capacity to enrich Zip3 at recombination initiation sites. Chromatin immunoprecipitation (ChIP) followed by qPCR to monitor the association of Zip3-FLAG with three DSB sites (*GAT1, BUΔ23, ERG1*, blue and green lines) as well as to centromere and axis sites (magenta and red lines, respectively) in SK1 strains carrying *ZIP1*, *zip1[Δ2-9]*, *zip1[Δ10-14]*, or *zip1Δ* alleles. *x* axes indicate number of hours after placement in sporulation medium. In the top graphs, the relative enrichment of indicated chromosomal sites detected by qPCR in Zip3-FLAG immunoprecipitates is expressed as a percentage of the level of each site detected in the input. In the lower graphs, the enrichment of indicated chromosomal sites detected by qPCR in Zip3-FLAG immunoprecipitates is normalized to the level of a negative control locus (internal to the 3.5 kb *NFT1* ORF) detected in Zip3-FLAG immunoprecipitates. Values are the average ± standard deviation from two independent experiments. A single experiment was performed in the case of the *zip1* null strain.

Similar to a *zip1* null mutant, little Zip3 was detectable at any of the three DSB sites examined, nor at the axis or centromere sites, in *zip1[Δ2-9]* or *zip1[Δ10-14]* strains (Figure 6). The phenotype of *zip1[Δ2-9]* in this experiment appeared indistinguishable from the *zip1* null, whereas slight Zip3 enrichment was detected at the *BUΔ23* DSB site in the *zip1[Δ10-14]* meiotic time course. These data indicate that the capacity of Zip1 to recruit Zip3 to DSB sites during meiosis relies on Zip1’s N terminal residues 2-14.

As expected based on ultrastructural images of recombination nodules along the length of synapsed chromosomes [20], using structured illumination microscopy (SIM) in conjunction with antibodies targeting the meiotic axis protein Red1 and targeting the central element protein(s) Ecm11 or Gmc2, we find that MutSγ and Zip3 foci localize at the midline of the SC, embedded within the SC central element: Singular foci of epitope-tagged MutSγ protein Msh4-MYC and Zip3-MYC localize directly between aligned Red1-labeled axes, where the SC central element substructure is positioned (Figures S6, S7). When antibodies targeting SC central element proteins are used to label the SC central element directly, Msh4 foci are embedded directly in the linear Ecm11-Gmc2 structures at the midline of the SC (Figure S7).

In SCs assembled by Zip1[Δ2-9] protein, we observe Zip3-MYC foci embedded within the SC central element, but such Zip3-MYC foci appear diminished in both number and intensity relative to Zip3-MYC foci on meiotic chromosomes from *ZIP1* strains (Figure S6). Similarly, using conventional wide-field fluorescence microscopy we observe a diminished number of bright Msh4-MYC foci on aligned mid–meiotic prophase chromosomes in strains expressing *zip1[Δ2-9]*, *zip1[Δ10-14]* and *zip1[Δ15-20]* relative to *ZIP1* strains (Figure S8); this phenotype mimics the Msh4-MYC pattern seen in a *zip1* null strain (Figure S8). Furthermore, the Msh4-MYC foci observed on synapsed meiotic prophase chromosomes in *zip1[Δ2-9]* strains do not robustly co-localize with other SIC proteins, such as Zip4-HA, as they do in wild-type synapsed nuclei (Figure S9).

Taken together, our ChIP and cytological studies indicate that residues within Zip1’s N terminal twenty amino acids are essential for the proper enrichment of the pro-crossover protein Zip3 to meiotic centromeres, to DSBs, to chromosomal axis sites, and for the accumulation of robust Zip3 and the MutSγ foci within the central region of the SC.

## Discussion

SC transverse filament proteins from different organisms share no ancestral homology but do share a conserved dual functionality: an activity that facilitates interhomolog crossover recombination events between DNA duplexes and a capacity to assemble the tripartite SC structure on meiotic chromosomes. Despite the absence of primary sequence conservation, SC transverse filament proteins from different phyla typically consist of an extended central “core” predicted to assemble coiled-coil, flanked by predicted unstructured N and C terminal regions. Sequence alignment of SC transverse filament proteins from related species reveal highly conserved residues throughout the central helical core of the protein, but also small groups of conserved amino acid residues within the N and C predicted unstructured domains (for a mammalian SC transverse filament example, see [47]; Figure 7A illustrates this point for the budding yeast transverse filament, Zip1). The small regions of conservation likely reflect functional residues, perhaps ones that form an interaction interface for a partner protein.

**fig 7.**
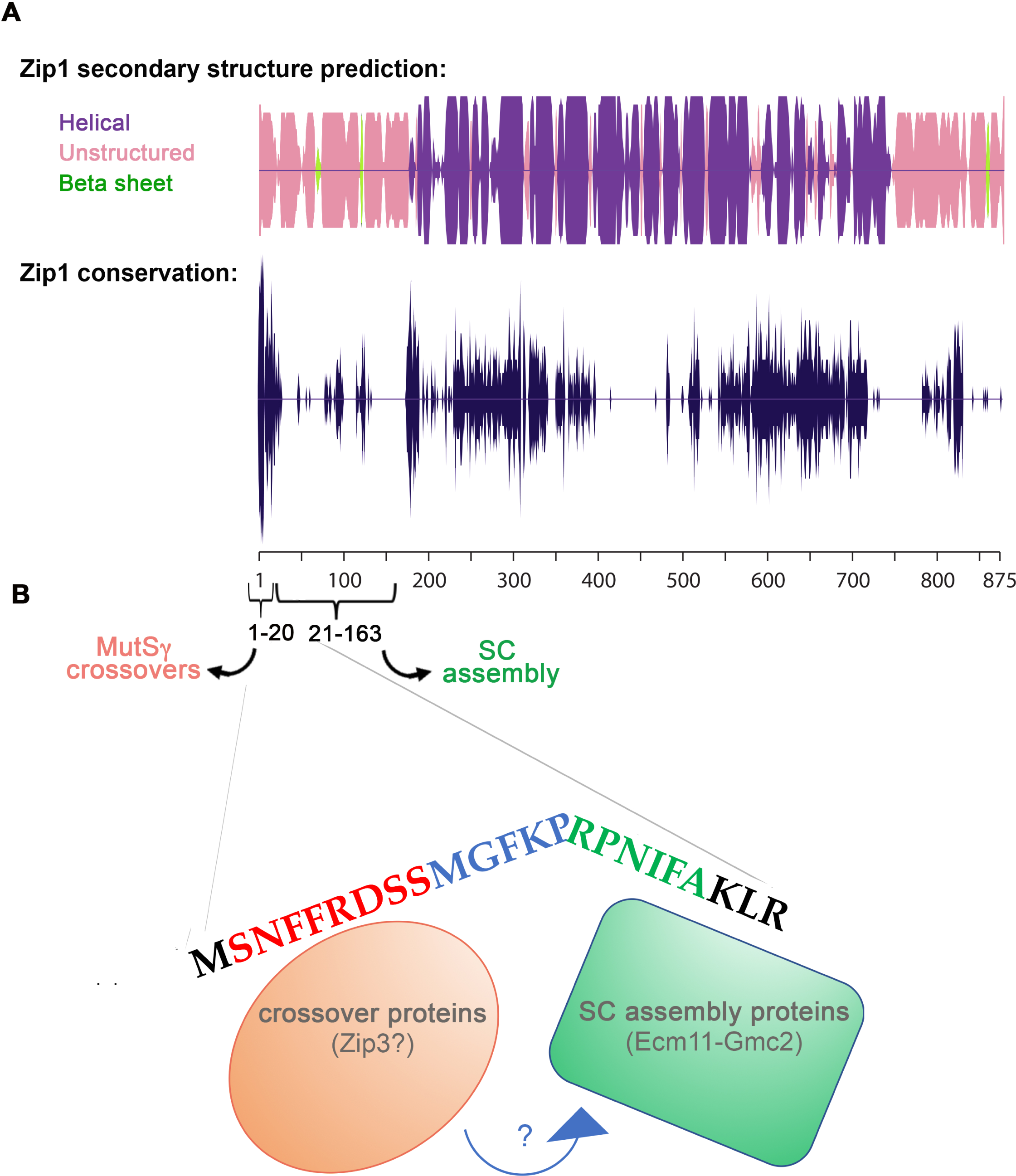
Adjacent conserved regions within Zip1’s N terminus coordinate meiotic crossing over with synapsis: A model. **A)** Cartoon illustrates the likelihood of the indicated secondary structure (alpha helical = purple; beta sheet = green; unstructured = pink) across the length of the 875 residue Zip1 protein. Secondary structure prediction was performed using JNet (http://www.compbio.dundee.ac.uk/www-jpred/). The lower line plot indicates the relative conservation of individual amino acid residues across the entire Zip1 protein, using homologs from the following fungal species: *Saccharomyces cerevisiae, Candida glabrata, Lachancea lanzarotensis, Tetrapisispora blattae, Tetrapisispora phaffii, Kazachstania naganishii, Vanderwaltozyma polyspora, Torulaspora delbrueckii, Zygosaccharomyces bailii, Zygosaccharomyces rouxii, Kazachstania africana, Naumovozyma dairenensis, Naumovozyma castellii, Kluyveromyces marxianus, Kluyveromyces dobzhanskii. Eremothecium cymbalariae, Ashbya gossypii, Ashbya aceri*. Stronger conservation is indicated by a longer line above and below the axis. Plots were created using Jalview 2.10 [58] and Adobe Illustrator CS5. The cartoon in **(B)**illustrates the possibility that Zip1’s N terminal 20 residues represents two regions with different functionalities: The region corresponding to residues 2-14 is required for MutSγ crossing over, to recruit the crossover regulator (and putative E3 SUMO ligase) Zip3 to meiotic DSB sites, and for the proper localization of Zip3 at Zip1 polycomplex structures. Our prior characterization of the *zip1[Δ21-163]* internal deletion allele indicates that residues between amino acids 21 and 163 are essential for Zip1’s SC assembly function but completely dispensable for MutSγ crossing over [22], in contrast to the reciprocal phenotype exhibited by *zip1[Δ2-9]* or *zip1[Δ10-14]*. We note that the region directly adjacent to residues 2-14 (corresponding to residues 15-20) is functionally important for both Zip1’s crossing over and synapsis activities, although this region has a bigger impact on Zip1’s SC assembly and Ecm11 SUMOylation activity (as demonstrated by the *zip1* null phenocopy displayed by *zip1[Δ15-20]* and *zip1[I18A, F19A]* strains) and plays a lesser role in Zip1’s pro-MutSγ crossover function relative to the 2-14 region. We speculate that the adjacency between these functionally distinct regions of Zip1’s N terminus may mechanistically underlie the coordination between MutSγ crossing over and synapsis, via direct molecular communication (blue arrow) between crossover factors and synapsis proteins.

In prior work we reported our discovery that a large in-frame deletion of Zip1’s predicted unstructured N terminal region (*zip1[Δ21-163];* originally created in [48]), encodes a separation–of–function Zip1 protein that fails to assemble mature SC but is completely capable of executing Zip1’s role in MutSγ crossing over [22]. Here we describe two novel non-null *zip1* alleles, *zip1[Δ2-9]* and *zip1[Δ10-14],* which confer the reciprocal separation-of-function phenotype: robust SC assembly in the absence of MutSγ crossovers. Together with our observation that *zip1[Δ2-163]* and *zip1[Δ2-20]* mutants abolish both SC assembly and MutSγ crossing over, these findings indicate that distinct regions within Zip1’s first twenty amino acid residues together serve both of Zip1’s separable functions. Consistent with this possibility, the Zip1’s first twenty residues represent the most highly conserved sequence in fungal Zip1 proteins (Figure 7A).

## Two functionally distinct regions within Zip1’s N terminal twenty residues: Interaction domains?

The analysis of our novel *zip1* alleles moreover leads us to propose the Zip1’s first twenty residues correspond to two distinct interaction domains, which directly engage with pro–crossover and pro–synapsis machinery and/or mechanisms. We propose that these pro-crossover and pro-synapsis domains are spatially positioned adjacent to one another within the N terminal tip of Zip1, as illustrated in the model shown in Figure 7B. One basis for our speculation is that Zip1 protein missing residues 2-9 or 10-14 confers a phenotype that strongly resembles the unique phenotype of cells missing the pro-crossover SIC protein, Zip3: *zip1[Δ2-9], zip1[Δ10-14]* and *zip3* strains [26, 28, 29] display SC assembly in the absence of MutSγ crossovers, and hyperSUMOylated forms of the SC central element protein, Ecm11. This phenotype contrasts with the synapsis-deficiency and diminished Ecm11 SUMOylation displayed by mutants missing Zip1 altogether, or missing other SIC proteins such as Zip2, Zip4, Spo16 or the MutSγ complex, which colocalize with Zip3 on meiotic chromosomes [3, 27, 29-31, 49], at recombination intermediates that are embedded in the central region of the SC (this work). The unique, *zip3*-like phenotype of *zip1[Δ2-9]* and *zip1[Δ10-14]* mutants suggests that these mutant strains are missing a specialized capacity to promote Zip3 function. (Although we note that synapsis is significantly more robust in *zip1[Δ2-9]* and *zip1[Δ10-14]* mutants, suggesting that Zip3 carries out additional pro–synapsis functions that are independent of Zip1’s N terminal tip residues.)

Additional support for the possibility that Zip1’s N terminal residues interact with Zip3 comes from the localization of Zip3 at polycomplex structures, aggregates of Zip1 and other SC-associated proteins that form when SC assembly is compromised. While many if not all SIC proteins have been found to localize to Zip1 polycomplex structures, Zip3’s localization shows a greater degree of coincidence with Zip1 throughout the bulk of the polycomplex, relative for example to Zip4 (Figure 5) or the MutSγ component, Msh4 [50], and this localization is completely abolished by the loss of Zip1’s residues 2-9 or 10-14 (Figure 5). Finally, Zip1[Δ2-9] and zip1[Δ10-14] have lost Zip1’s capacity to recruit Zip3 to sites of recombination initiation (Figure 6).

Based on the absence of SC assembly in meiotic cells expressing *zip1[Δ15-20]*, *zip1[15-20→A]* and *zip1[I18A, I19A]*, we furthermore conclude that Zip1’s first twenty residues correspond to (perhaps in an overlapping manner) at least one interaction domain for an SC assembly factor or complex of factors. Unlike the limited nature of the region within Zip1’s N terminal 163 amino acids that is required for MutSγ crossovers (twenty residues, based on the fact that *zip1[Δ21-163]* is fully capable of MutSγ crossing over [22]), groups of residues that are critical for allowing Zip1 to assemble SC may be distributed throughout the entire N terminal region encompassed by residues 15-163, as both *zip1[Δ15-20]* and *zip1[Δ21-163]* fail to assemble SC. Nevertheless, we propose that the region overlapping residues 15-20 interfaces with components serving an SC assembly function, based on the fact that alteration of two adjacent residues (I18 and I19) at Zip1’s extreme N terminus (a change that is unlikely to alter the overall length or structure of the rod-like protein) completely abolishes Zip1’s capacity to assemble SC.

We previously demonstrated that *zip1[Δ21-163]* phenocopies the *ecm11* and *gmc2* null mutant phenotype (a failure in SC assembly but proficiency in MutSγ crossing over), suggesting that this N terminal region of Zip1 functionally interacts with the central element in order to assemble SC [22]. Moreover, Leung et. al (2015) demonstrated that Zip1’s N terminal 346 residues is sufficient to promote Ecm11 SUMOylation in vegetative (non-meiotic) cells, provided that Gmc2 is also expressed. These data suggest that the N terminal region of Zip1 is able to engage with the Ecm11-Gmc2 proteins, perhaps in a direct manner or perhaps indirectly through a protein expressed in both meiotic as well as mitotic cells [51]. Similar to the uncertainty about whether Zip3 interacts with Zip1 in a direct manner, apart from the genetic interactions found for Zip1 and Ecm11 and the coincidence of Ecm11 and Gmc2 at the midline of SC (where Zip1 N termini also reside [19]), strong evidence of a direct physical interaction between Zip1 and Ecm11 or Gmc2 does not yet exist.

## The adjacency between Zip1’s pro-crossover and a pro–synapsis regions may serve as a liaison to coordinate SC assembly with intermediate steps in the MutSγ crossover pathway

Finally, we note the tantalizing possibility that the *adjacency* between the putative pro-crossover and pro-synapsis regions of Zip1’s N terminus is functionally important for ensuring that SC assembly occurs in coordination with intermediate steps in the MutSγ crossover recombination pathway. Specifically, we speculate that Zip1 may physically connect crossover recombination events to SC assembly through a mechanism that is based, at least in part, on its capacity to stabilize Zip3 at its N terminus. Here, Zip3 (which has SUMO transferase activity [40]) would be expected to be in close proximity to putative pro–synapsis factors stabilized by the adjacent region in Zip1, and could perhaps regulate the extent of SUMOylation of SC central element protein Ecm11. Perhaps negative regulation of Ecm11 SUMOylation by Zip3 is directly involved in ensuring that SC assembly occurs “in the right place at the right time” (i.e. at a MutSγ crossover event).

## SCs assembled in the absence of MutSγ crossing over in budding yeast may be less stable

During the course of analyzing SC assembly over a time course of meiotic progression in our mutants we obtained an unexpected finding: SCs assembled in *zip3, zip1[Δ2-9]* and *zip1[10-14]* strains (corresponding to SCs assembled in the absence of MutSγ crossovers) assemble earlier than wild-type SC structures, and appear to be less capable of persisting during an *ndt80-* mediated, meiotic prophase arrest (Figure 3). Pattabiraman (2017) found that a MutSγ-associated process affects the dynamic properties of *C. elegans* SC [33]; our set of preliminary observations raises the intriguing possibility that the MutSγ crossover pathway influences the structure and/or dynamics of budding yeast SCs in a similar fashion.

## Methods

### Strains

Yeast strains used in this study are isogenic to BR1919-8B [52] and were created using standard genetic crosses and manipulation procedures. CRISPR-Cas9 methodology was utilized to create unmarked alleles. Crossover analysis strains carry an *hphMX4* cassette inserted near the chromosome III centromere, *ADE2* inserted upstream of the *RAD18* locus, a *natMX4* cassette inserted near the *HMR* locus, *TRP1MX4* was inserted 62 bp downstream of the *SPO11* locus (Kee and Keeney, 2002), *URA3* replacing *SPO13,* and *LYS2* inserted on chromosome VIII at coordinate 210,400 bp. Strains SYC107, SYC149, SYC151 and K914 do not carry *THR1* nor *LYS2* on chromosome VIII. ChIP and qPCR experiments were performed in strains of the SK1 genetic background (Table S4).

### Cytological Analysis and Imaging

Meiotic nuclei were surface spread on glass slides and imaged as described in [22]. The following primary antibodies were used: affinity purified rabbit anti-Zip1 (1:100, raised at YenZym Antibodies, LLC, against a C terminal fragment of Zip1, as described in [38], mouse anti-cMYC (1:200, clone 9E10, Abcam). Mouse anti-Gmc2 antibodies were raised againsed purified Gmc2 protein, and guinea pig anti-Gmc2_Ecm11 antibodies were raised against a co-purified protein complex (ProSci Inc.). These antibodies were used at 1:800. Chicken anti-HA (1:100, Abcam), and rabbit anti-Red1 (1:100, a kind gift from G.S. Roeder, [53]) were also used. Secondary antibodies conjugated with Alexa Fluor dyes were purchased from Jackson ImmunoResearch and used at 1:200 dilution. Microscopy and image processing was carried out using a Deltavision RT imaging system (Applied Precision) adapted to an Olympus (IX71) microscope. Structured illumination microscopy was carried out using Applied Precision’s OMX Blaze Structured Illumination Microscope system at The Rockefeller University’s Bio-Imaging Resource Center.

### Calculations and Statistical Analysis

Genetic crossover data was compiled and processed using an Excel Linkage Macro program, created by Jonathan Greene (Rhona Borts, pers. comm.) and donated by Eva Hoffmann (University of Copenhagen, Denmark). Crossover values (and their standard errors) were obtained using the Stahl lab online tools (http://molbio.uoregon.edu/∼fstahl/), with the method of Perkins [54]. Non-mendelian segregation is reported in Table S3. Recombinant spore values were calculated according to the following: 100(*r/t*), where *r* = the number of colonies carrying a chromosome which is recombinant in the interval and *t*; the total number of colonies assessed. Standard error (S.E.) values for random spore analysis were calculated according to the formula: 100(√(r/t)(1-r/t)/t) [55]. All other statistical analyses were carried out using Graphpad Prism or Graphpad InStat (www.graphpad.com).

### Western Blot

Western blotting was performed as described previously [19] with the following modifications: Amersham Protran 0.2μm NC was used as the transfer membrane following the manufacturer’s recommendation; after secondary antibody incubation the membrane was processed with a final wash in 100mM Tris-Cl pH 9.5, 100mM NaCl, 5mM MgCl_2_ to boost the HRP-mediated chemiluminescence using Amersham ECL Prime Western Blotting Detection Reagent.

### Chromatin Immunoprecipitation

Meiotic cells were processed as described [56], with the following modifications: Lysis was performed in Lysis buffer plus 1 mM PMSF, 50 µg/mL Aprotinin and 1X Complete Mini EDTA-Free (Roche), using 0.5 mm zirconium/silica beads (Biospec Products, Bartlesville, OK). 2 µg of the mouse monoclonal anti-FLAG antibody M2 (Sigma) and 30 µL Protein G magnetic beads (New England Biolabs) were used. Quantitative PCR was performed from the immunoprecipitated DNA or the whole–cell extract using a 7900HT Fast Real-Time PCR System (Applied Biosystems, Thermo Scientific) and SYBR Green PCR master mix (Applied Biosystems) as described [56]. Results were expressed as % of DNA in the total input present in the immunoprecipitated sample and/or normalized to the negative control site in the middle of *NFT1*, a 3.5 kb long gene. Primers for *GAT1*, *BUΔ23*, *ERG1*, Axis and *NFT1* loci have been described [45, 46, 57].

## Supplemental Information

**S1 Fig.**
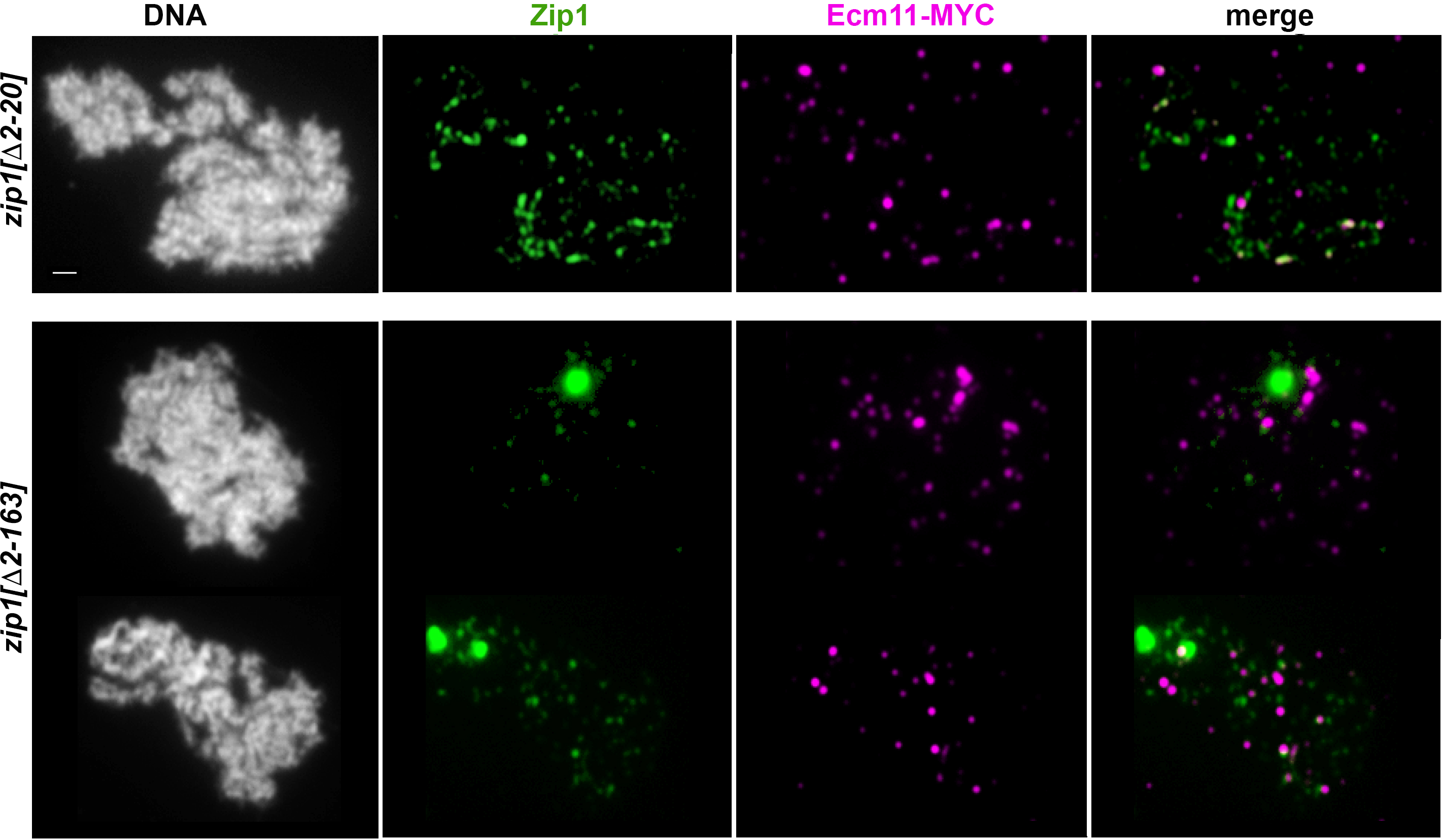
Residues within Zip1’s 2-20 region are required for SC assembly. Representative surface-spread mid-meiotic prophase nuclei from diploids homozygous for the *zip1[Δ2-20]* (top row), or *zip1[Δ2-163]* (bottom row) alleles. (Genotypes indicated at left.) All strains also carry the *ndt80* null allele, which allows meiotic cultures to accumulate at mid-late prophase stages when full-length SCs are normally present. Mid-meiotic prophase chromosomes are stained with DAPI to label DNA (white), anti-Zip1 to label Zip1 (green), and anti-MYC to label Ecm11 (magenta). The merge between Zip1 and Ecm11 channels is shown in the final column. Scale bar, 1 µm.

**S2 Fig.**
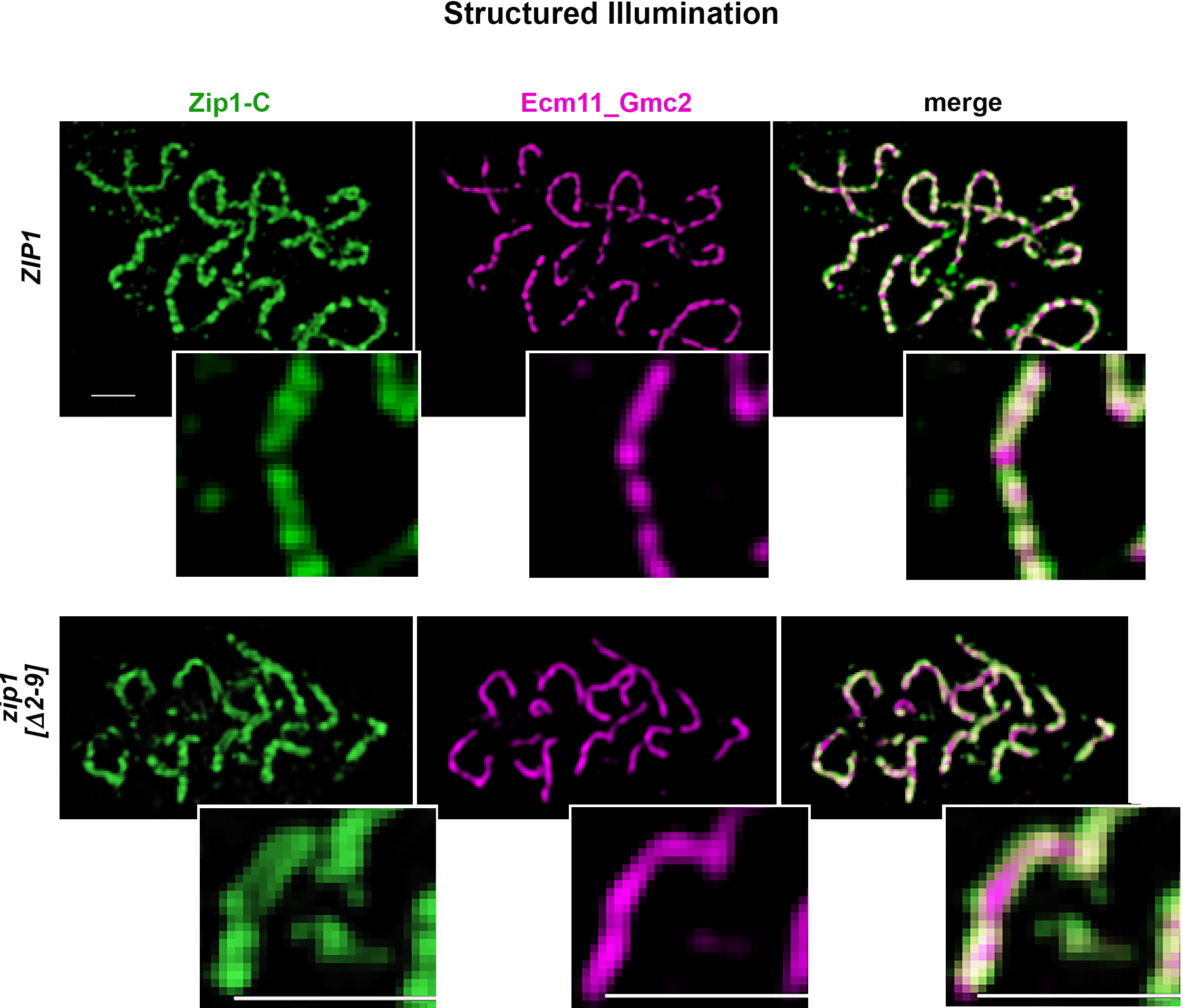
Zip1[Δ2-9] assembles with the proper orientation within tripartite SC. Structured illumination was used to probe the organization Zip1 and Ecm11 within linear assemblies in wild type (top panels) and *zip1[Δ2-9]* (bottom panels) strains. Antibodies against the C terminal 264 residues of the Zip1 protein (green) were applied in conjunction with antibodies against a mixture of Ecm11 and Gmc2 peptides (magenta; see Methods for antibody information). In surface-spread meiotic nuclei containing wild-type SC (top panels), anti-Zip1 is often detected as a wide ribbon in which parallel tracts are sometimes visible with the resolution power of structured illumination (∼120 nm). Such parallel tracts of Zip1 (see zoomed insets) flank a central narrow band of Ecm11-Gmc2 protein, and reflect Zip1’s orientation within the mature SC, where Zip1’s C termini interface with lengthwise-aligned homologous chromosome axes and N termini interface closer to the SC central element (comprised of Ecm11 and Gmc2). DNA is labeled by DAPI (white). Bottom panels show a representative spread where wild-type Zip1 organization is seen within the SCs built by the Zip1[Δ2-9] protein. Scale bar, 1 μm.

**S3 Fig.**
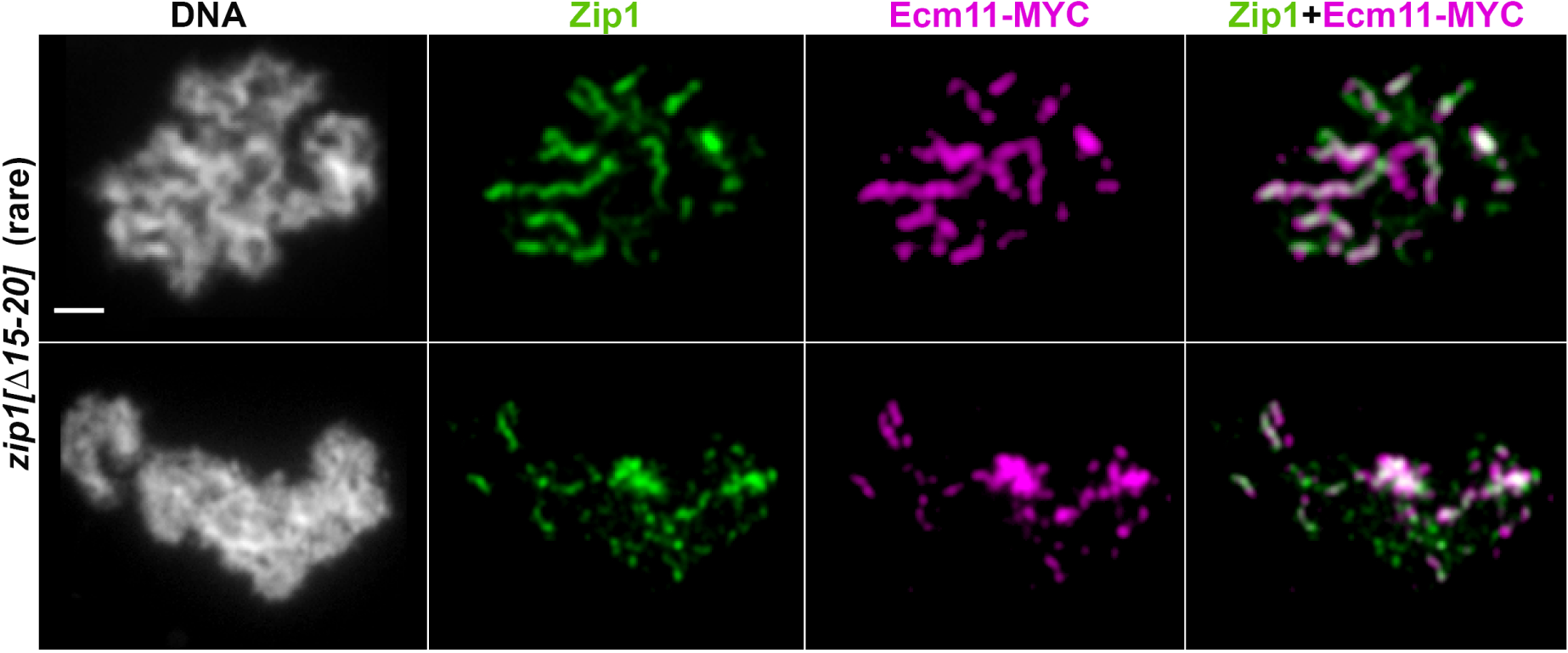
Zip1[Δ15-20] occasionally assembles fragile coincident Zip1 and Ecm11 structures. While the vast majority of meiotic nuclei from *zip1[Δ15-20]* strains exhibited only abundant and varying-sized foci of Zip1 and Ecm11, occasionally frail-looking linear assemblies of diffuse Zip1 (green) would accompany similar types of Ecm11 linear assemblies (magenta) on surface-spread meiotic chromosomes (labeled with DAPI, white). These assemblies (as well as instances of adjacent large focal deposits of Zip1 and Ecm11) were included in our linear assembly measurements (Fig. 3). The abnormal-looking linear assemblies appear wavy, and often taper at their ends. The top row presents the one rare nucleus out of the (more than 100) nuclei examined, in which these frail linear assemblies were most abundant. We note that unlike the robust linear assemblies of coincident Ecm11 and Zip1 observed in *zip1[Δ2-9]* or *zip1[Δ10-14]* strains (Fig.2), these diffuse linear assemblies do not appear to join lengthwise-aligned chromosomes. Scale bar, 1 mm.

**S4 Fig.**
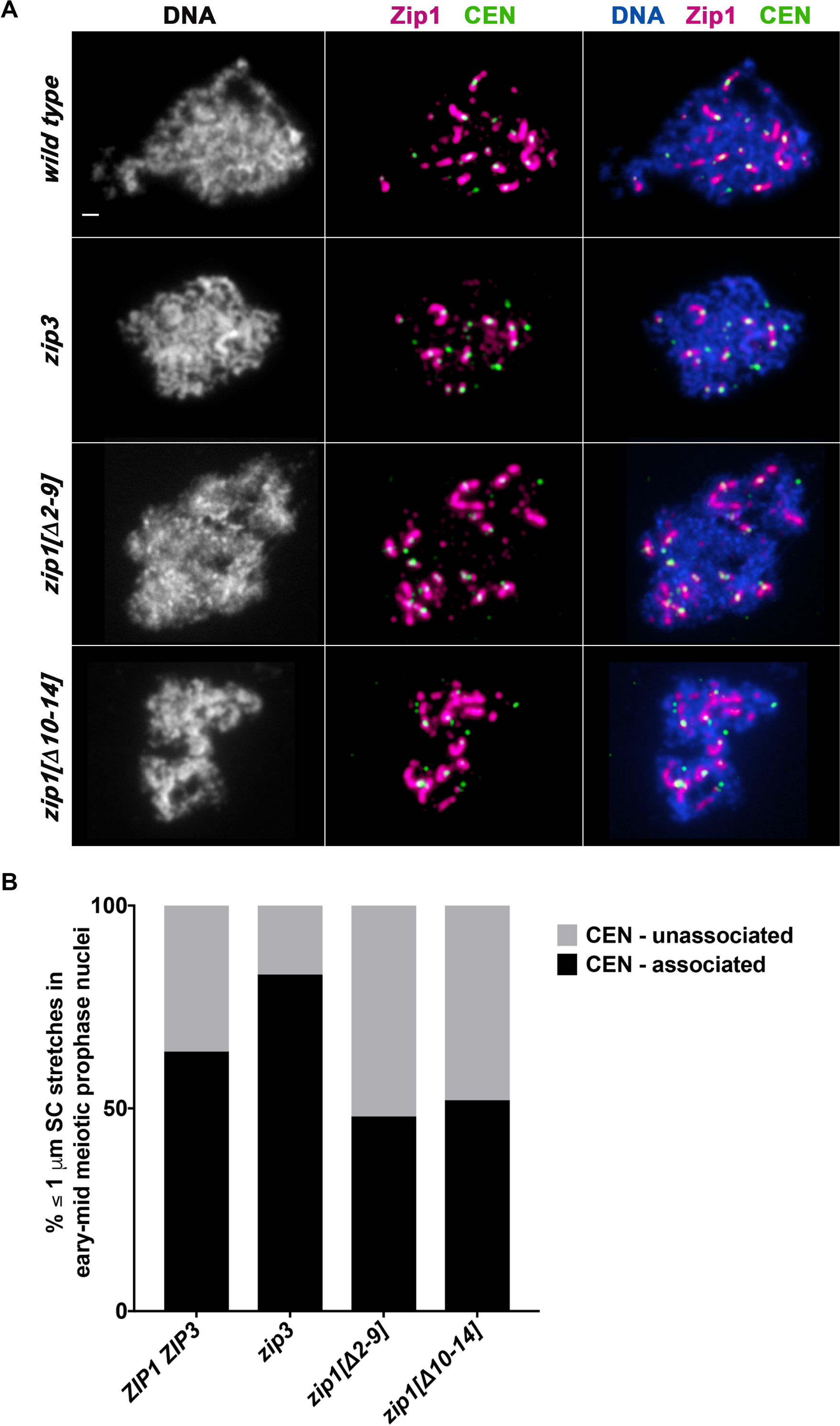
SC comprised of Zip1[Δ2-9] or Zip1[Δ10-14] initiates at both centromeric and noncentromeric chromosomal sites. **A)**Representative surface-spread meiotic nuclei from wild-type (top row), *zip3* (second row), *zip1[Δ2-9]* (third row) or *zip1[Δ10-14]*, with DAPI-labeled DNA (white), anti-Zip1 (magenta), and anti-MYC to label the centromere-associated Ctf19-MYC protein (green). While all strains at this early stage of synapsis exhibited a large number of short Zip1 assemblies that were associated with a centromere, fewer examples of ≤1 μm Zip1 assemblies without an associated centromere were observed in *zip3* strains, relative to wild-type, *zip1[Δ2-9]* or *zip1[Δ10-14]* strains. Scale bar, 1 μm. Quantitation of the number of ≤1 μm Zip1 assemblies with or without an associated Ctf19-MYC signal is shown in the bar graphs in **(B)**, where light shading represents the percentage of total ≤1 μm Zip1 assemblies in each strain that are unassociated with a centromere signal. In wild-type strains, 23 out of 36 ≤1 µm Zip1 assemblies (in 10 surface-spread nuclei) were associated with a centromere; In *zip3* strains, 29 out of 35 ≤1 µm Zip1 assemblies (in 11 nuclei) were associated with a centromere; In *zip1[Δ2-9]* strains, 35 out of 73 ≤1 μm Zip1 assemblies (in 21 nuclei) were associated with a centromere; In *zip1[Δ10-14]* strains, 27 out of 52 ≤1 μm Zip1 assemblies (in 11 nuclei) were associated with a centromere. A Fishers Exact Test found no significant difference between the proportion of centromere-associated Zip1 stretches in wild-type versus *zip3* in this data set (two-tailed *P* value = 0.107), but did find a significant difference between *zip1[Δ2-9],* and *zip3* (two-tailed *P* value =0.0007), and between *zip1[Δ10-14]* and *zip3* (two-tailed *P* value = 0.0033).

**S5 Fig.**
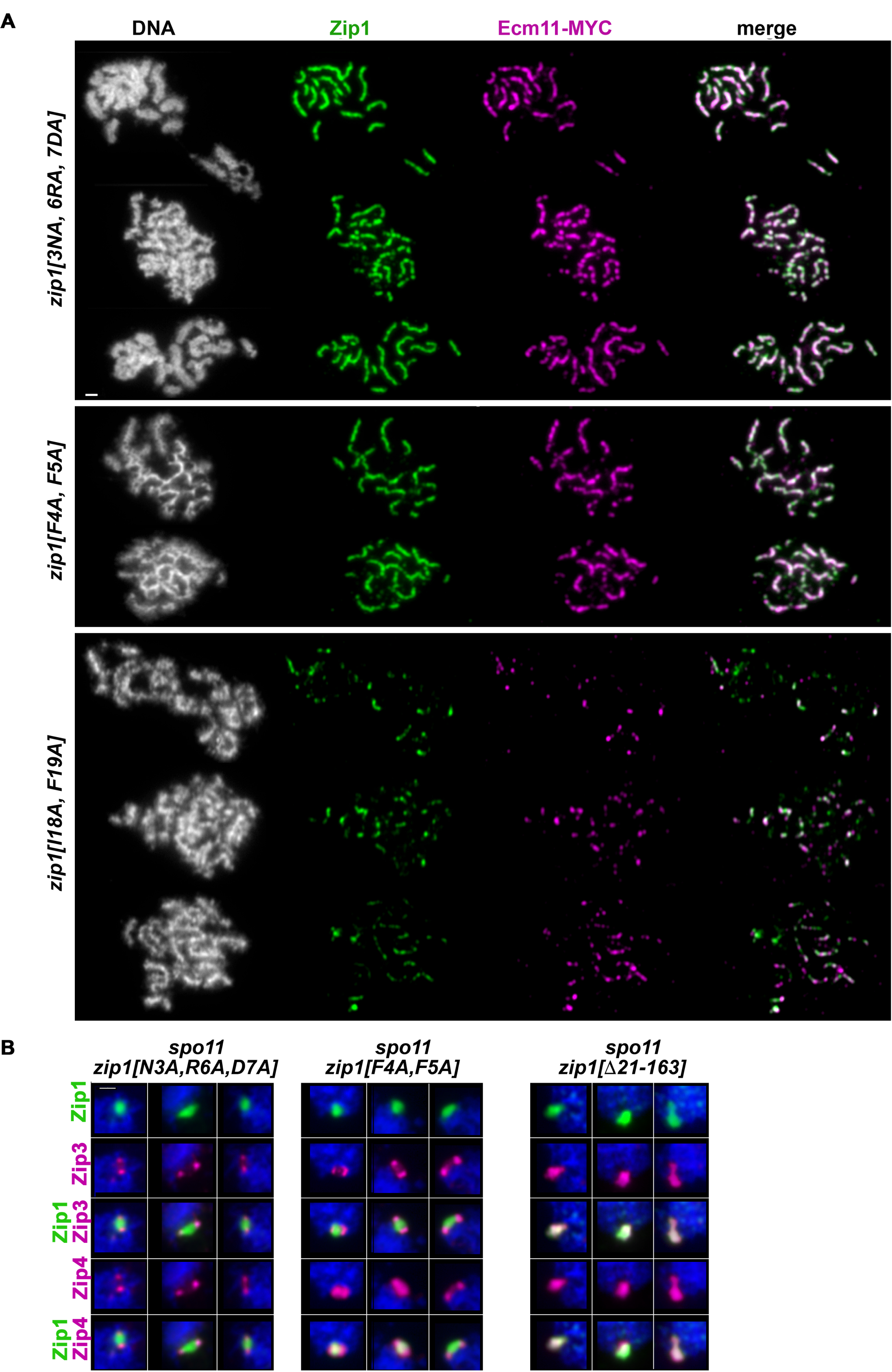
*zip1[N3A, R6A, D7A]* and *zip1[F4A, F5A]* and *zip1[I18A, F19A]* mutants resemble corresponding small deletion alleles with respect to synapsis and Zip3 localization to polycomplex. **A)** Panels show representative surface-spread mid-meiotic prophase nuclei from *zip1[N3A, R6A, D7A]* (top panel, including 3 rows), *zip1[F4A, F5A]* (middle panel, including two rows) and *zip1[I18A, F19A]* (bottom panel, including three rows) mutants (genotypes indicated at left). All strains carry an *ndt80* null allele, which allows meiotic cultures to accumulate at mid-late prophase stages when full-length SCs are normally present. Mid-meiotic prophase chromosomes are stained with DAPI to label DNA (white), anti-Zip1 to label Zip1 (green), and anti-MYC to label epitope-tagged Ecm11 (magenta). The merge between Zip1 and Ecm11 channels is shown in the final column. (**B**) Three groups of panels correspond to three representative images of Zip1 polycomplex structures (first row, green) in *spo11* meiotic prophase nuclei homozygous for either *zip1[N3A, R6A, D7A]* (far left group), *zip1[F4A, F5A]* (middle group), or *zip1[Δ21-163]* (far right group). The localization of Zip3-MYC (second row, magenta) and Zip4-HA (fourth row, magenta) is assessed on these surface-spread meiotic prophase nuclei. Merged images between either Zip3-MYC and Zip1 or Zip4-HA and Zip1 are shown in the third and fifth rows, respectively. Scale bar, 1 μm.

**S6 Fig.**
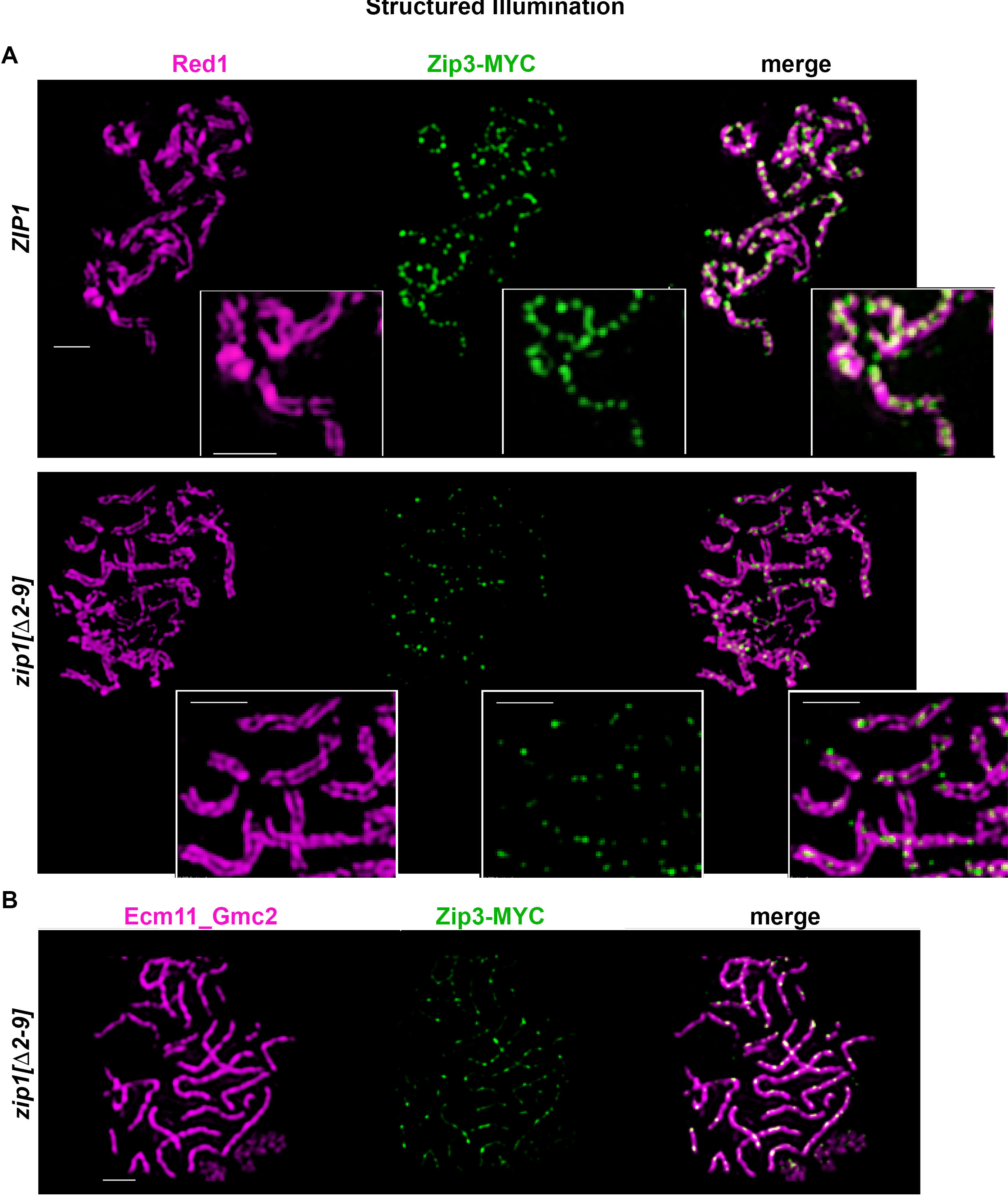
Zip3-MYC foci localize to the SC central element sub-structure in wild-type meiotic nuclei but are diminished in number and intensity in *zip1[Δ2-9]* and *zip1[Δ10-14]* strains. Images show representative surface-spread meiotic nuclei labeled with antibodies against the chromosomal axis protein Red1 (magenta) and antibodies against the MYC eptitope to label Zip3-MYC (green). In wild-type strains (top panels), structured illumination microscopy reveals Zip3-MYC foci directly in between lengthwise-aligned homologous chromosome axes in mid-meiotic prophase nuclei, reflecting the embedded distribution of Zip3 complexes within the central region of SCs. The position of Zip3-MYC foci in between lengthwise-aligned axes is unchanged but the number of robust Zip3-MYC foci is severely diminished in *zip1[Δ2-9]* strains (lower panels). Zip3-MYC’s location within the central element substructure of the SC in this strain is also indicated in the bottom row of images, where Zip3-MYC (green) is co-labeled with antibodies that target Ecm11-Gmc2 (magenta). Scale bar, 1 µm.

**S7 Fig.**
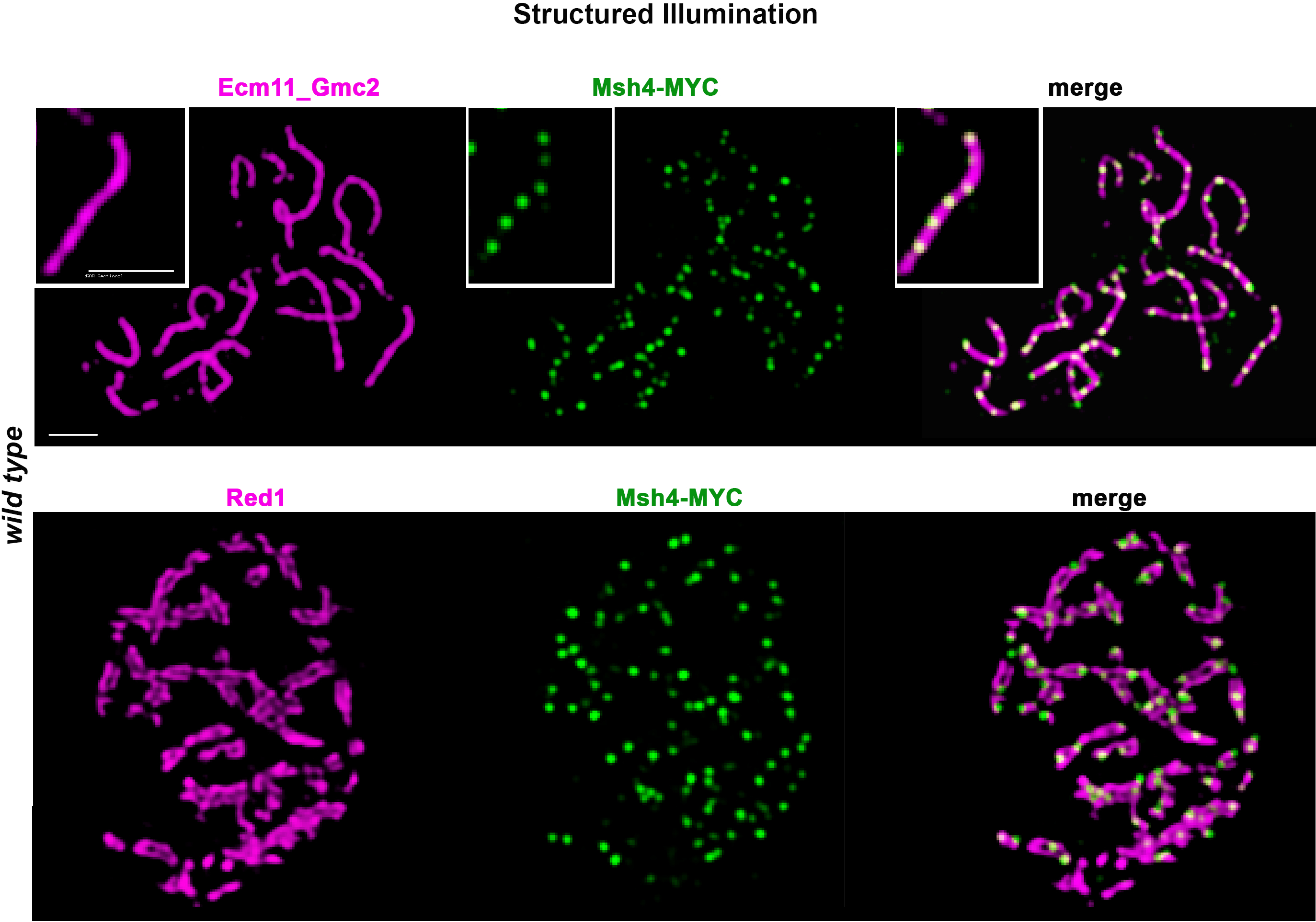
Msh4-MYC localizes to the SC central element sub-structure. Images show representative surface-spread meiotic nuclei from wild-type strains, labeled with antibodies against Ecm11-Gmc2 central element proteins (magenta, top row), or the chromosomal axis protein Red1 (magenta, bottom row) and antibodies against the MYC epitope to label a MYC-tagged version of the MutSγ component, Msh4 (green). The increased resolving power of structured illumination microscopy reveals that Msh4-MYC foci directly embed within the central element substructure of the SC (see zoomed inset in top row). Scale bar, 1 μm.

**S8 Fig.**
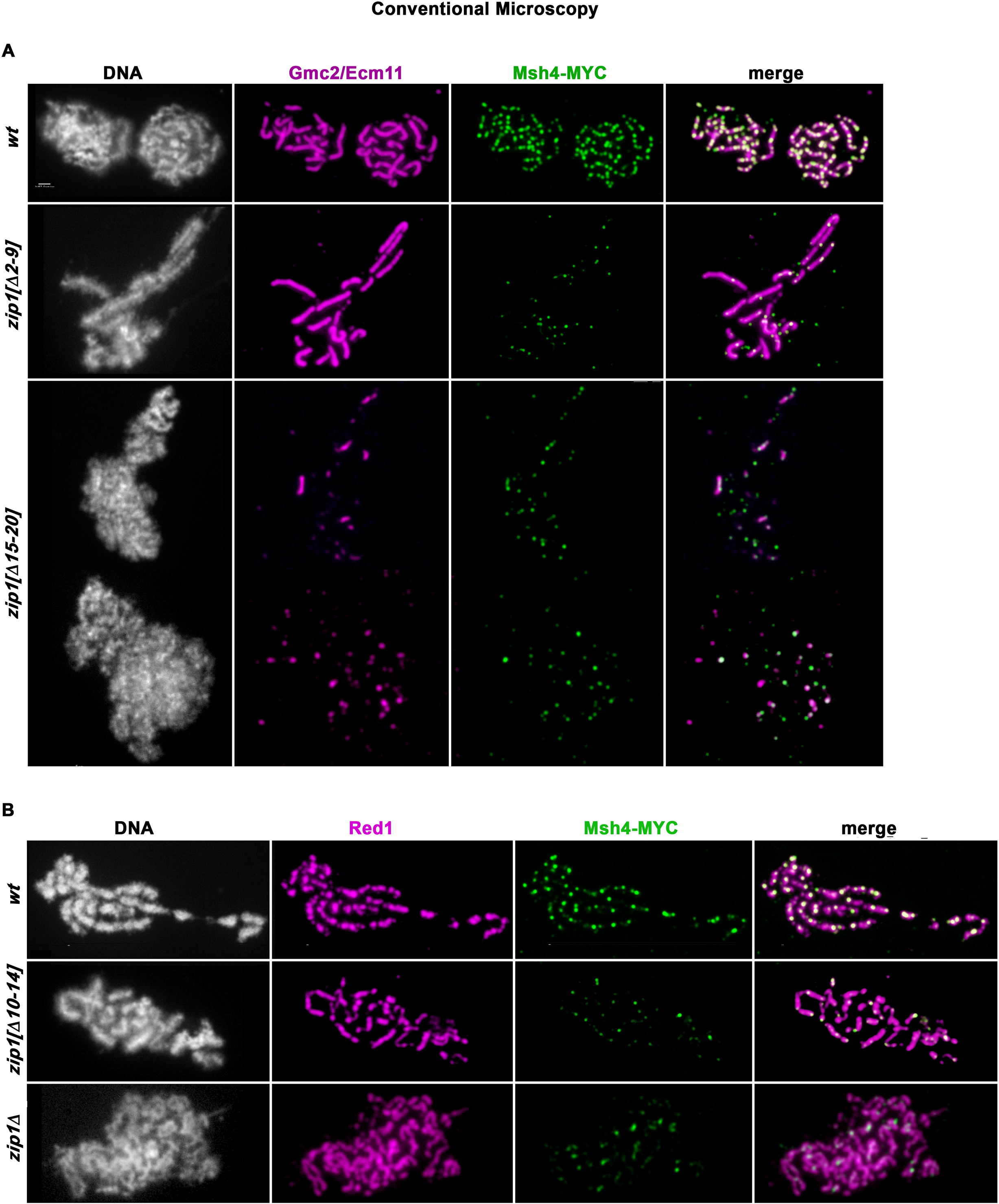
Msh4-MYC foci are reduced in number and diminished in size in *zip1[Δ2-9]* and *zip1[Δ10-14]* strains. Images show representative surface-spread meiotic nuclei labeled with antibodies that target Ecm11-Gmc2 (magenta, top rows) or antibodies against the chromosomal axis protein Red1 (magenta, bottom rows) in conjunction with antibodies that target the MYC epitope to label Msh4-MYC (green). (**A**) In wild-type strains (top row), conventional fluorescence microscopy reveals Msh4-MYC foci directly embedded within the SC central element substructure, consistent with a similar number of Zip3 complexes (Fig. S6), which colocalize with Msh4 [26], within the central region of SCs. The position of Msh4-MYC foci within the central region of the SC is unchanged but the number of robust Msh4-MYC foci is severely diminished in *zip1[Δ2-9]* strains (middle row) and in *zip1[Δ15-20]* strains (bottom panel including two rows). The distribution of Msh4-MYC foci with respect to Red1-labeled chromosome axes is shown in (**B**; top row), and these images reveal a similar diminishment in bright Msh4-MYC foci in *zip1[Δ10-14]* strains (middle row), as well as *zip1* null strains (bottom row). Scale bar, 1 μm.

**S9 Fig.**
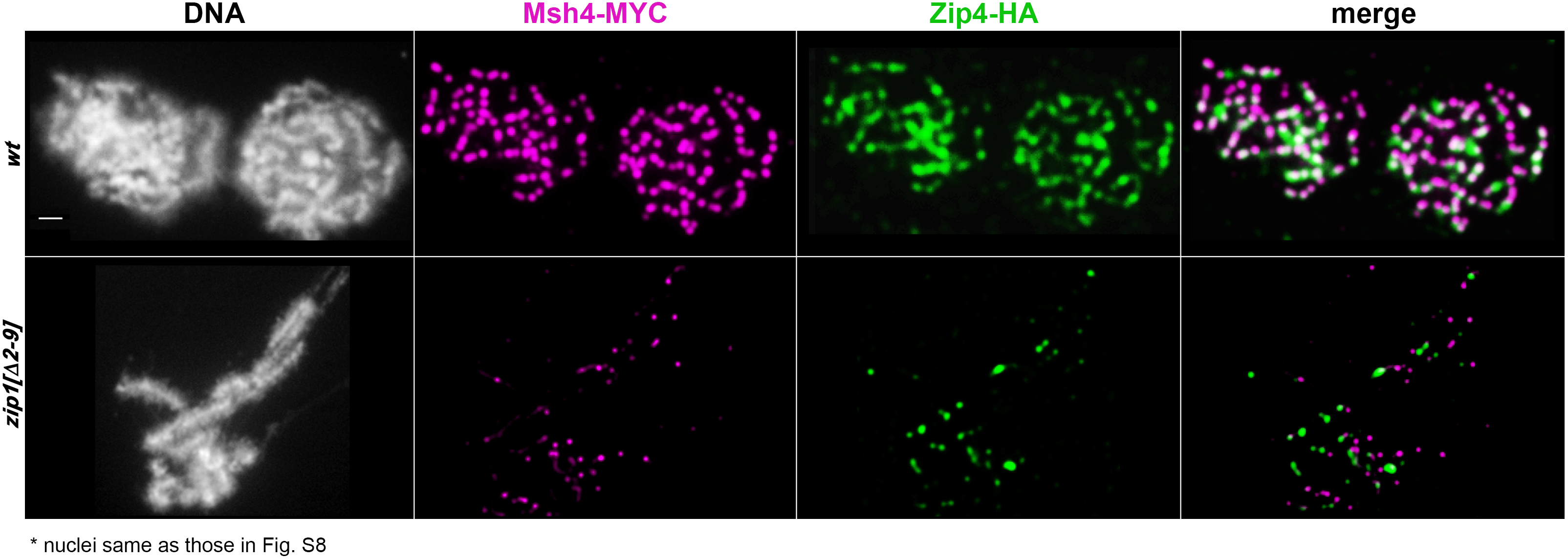
Msh4-MYC foci rarely co-localize with Zip4-HA in *zip1[Δ2-9]* strains. Images display the same surface-spread meiotic nuclei shown in the top two rows of Fig. S8. In addition to DNA (white) and Msh4-MYC (magenta), Zip4-HA is labeled in green. In wild-type strains (top row), Msh4-MYC foci generally appear coincident to or adjacent to Zip4-HA foci. By
contrast, the fewer and less bright Msh4-MYC and Zip4-HA foci that localize to synapsed meiotic prophase chromosomes in *zip1[Δ2-9]* strains rarely appear co-localized. Scale bar, 1 μm.

**S1 Table.** Sporulation efficiency and spore viability of strains used for crossover analysis. Sporulation efficiency reflects the fraction of cells that are 2, 3 or 4-spore asci after 5 days on sporulation plates. The frequency of tetrads containing four, three, two, one, or zero viable spores is shown along with the total spore viability (under “% Spore viability”); n.d.= not determined. Full strain genotypes are listed in Table S4. An asterisk indicates data that was previously published [22, 39].

|  |  | SPORULATION |  |  |  |  |  |  | % Spore<br>viability | Chromosome III<br>Non-Disjunction |  |
| --- | --- | --- | --- | --- | --- | --- | --- | --- | --- | --- | --- |
|  |  | EFFICIENCY<br>% (n) | # Tetrads | % 4 Spore<br>viable | % 3 Spore<br>viable | % 2 Spore<br>viable | % 1 Spore<br>viable | % 0 Spore<br>viable |  | (NDJ) % | #NDJ/<br># 2 spore viable |
| *pch2Δ* | (AM3724) | 76 (1100) | 757 | 73 | 17 | 8 | 1 | 1 | **90** | 3.3 | 2/61 |
| *pch2Δ msh4Δ* | (AM4025) | 12 (1031) | 826 | 25 | 13 | 23 | 17 | 22 | **51** | 10.1 | 10/189 |
| *pch2Δ zip1Δ* | (AM4023) | 15 (1023) | 828 | 13 | 14 | 21 | 20 | 31 | **39** | 0.6 | 1/175 |
| *pch2Δ zip1Δ msh4Δ* | (AM4026) | 14 (1037) | 453 | 6 | 7 | 22 | 27 | 39 | **29** | 7.1 | 7/98 |
| *pch2Δ zip1[Δ2-163]* | (AM3725) | 28 (1066) | 1098 | 31 | 16 | 22 | 15 | 16 | **58** | 4.2 | 10/238 |
| *zip1[Δ2-163]* | (AM3655) | 1 (1010) |  |  |  |  |  |  |  | n.d. |  |
| *zip1Δ \** | (CO10) | 5 (4503) | 80 | 28 | 23 | 20 | 6 | 23 | **56** | n.d. | ND |
| *WT\** | (K842) | 57 (1007) | 786 | 92 | 4 | 3 | 0 | 0 | **97** | 0.0 | 0/27 |
| *msh4Δ \** | (K852) | 42 (1005) | 1028 | 42 | 18 | 23 | 15 | 19 | **71** | 11.0 | 26/239 |
| *zip1[Δ2-20]* | (AM3684) | 20 (1061) | 1030 | 60 | 17 | 13 | 4 | 6 | **81** | 21.6 | 29/134 |
| *zip1[Δ2-20] msh4Δ* | (K1000) | 12 (1053) | 961 | 46 | 16 | 17 | 9 | 12 | **69** | 20.4 | 33/162 |
| *zip1[Δ2-9]* | (MP43) | 57 (1011) | 852 | 71 | 15 | 8 | 3 | 2 | **87** | 2.9 | 2/69 |
| *zip1[Δ2-9] msh4Δ* | (MP46) | 36 (1017) | 1287 | 38 | 17 | 21 | 12 | 12 | **64** | 4.8 | 13/269 |
| *zip1[Δ10-14]* | (SYC107) | 54 (3196) | 802 | 79 | 13 | 6 | 1 | 1 | **92** | 2.1 | 1/48 |
| *zip1[Δ10-14] msh4* | (SYC149) | 5 (502) | 750 | 50 | 17 | 15 | 10 | 8 | **73** | 8.6 | 10/116 |
| *zip1[Δ15-20]* | (AF8) | 51 (1044) | 802 | 72 | 12 | 11 | 2 | 2 | **87** | 14.4 | 13/90 |
| *zip1[Δ15-20] msh4Δ* | (K914) | 34 (1048) | 968 | 57 | 15 | 16 | 6 | 6 | **78** | 18.4 | 29/158 |
| *zip1[Δ21-163]* | (AF6) | 42 (1107) | 1025 | 60 | 22 | 12 | 4 | 1 | **84** | 0.8 | 1/128 |
| *zip1[Δ21-163] msh4Δ\** | (SYC151) | 41 (1243) | 1056 | 59 | 11 | 16 | 3 | 11 | **76** | 8.6 | 10/116 |
| *zip3Δ* | (K926) | n.d. | 866 | 61 | 16 | 14 | 5 | 5 | **81** | 10.0 | 12/120 |
| *zip3Δ msh4Δ* | (AM3658/AM3659) | 10 (2136) | 1049 | 58 | 14 | 18 | 4 | 6 | **79** | 5.4 | 10/185 |
| *zip3Δ zip1[Δ2-9]* | (MP52) | 49 (1014) | 1089 | 51 | 15 | 18 | 8 | 8 | **73** | 8.7 | 17/196 |
| *zip1[N3A,R6A,D7A]* | (K1281) | 61 (1134) | 660 | 81 | 12 | 6 | 1 | 1 | **93** | 5.4 | 2/37 |
| *zip1[F4A,F5A]* | (K1309) | 54 (1224) | 624 | 69 | 18 | 9 | 2 | 3 | **87** | 3.4 | 2/59 |
| *zip1[I18A,F19A]* | (K1282) | 54 (1235) | 268 | 81 | 9 | 4 | 1 | 3 | **91** | 8.3 | 1/12 |

\* indicates data that was previously published (Voelkel-Meiman, Johnston et al. 2015, Voelkel-Meiman 2016)

**S2 Table.** Map distances in *zip3* and additional *zip1* allele strains. Data display and calculations are as in Table 2.

| GENOTYPE<br>(STRAIN) | INTERVAL<br>(CHROMOSOME) | PD | TT | NPD | TOTAL | cM<br>(± SE) | %WT | cM<br>by chrM | %WT<br>by chrM | NPDobs/NPDexp<br>(± SE) |
| --- | --- | --- | --- | --- | --- | --- | --- | --- | --- | --- |
| *zip1*[N3A,R6A,D7A]<br>(K1281) | *HIS4-CEN3* (III) | 304 | 219 | 3 | 526 | **22.5 (1.4)** | **84** | 83.1 (III) | **78** | 0.18 (0.11) |
|  | *CEN3-MAT* (III) | 367 | 156 | 1 | 524 | **15.5 (1.1)** | **77** |  |  | 0.14 (0.14) |
|  | *MAT-RAD18* (III) | 264 | 255 | 5 | 524 | **27.2 (1.6)** | **75** |  |  | 0.20 (0.09) |
|  | *RAD18-HMR* (III) | 354 | 171 | 3 | 528 | **17.9 (1.4)** | **78** |  |  | 0.33 (0.19) |
|  | *SPO11-SPO13* (VIII) | 289 | 223 | 5 | 517 | **24.5 (1.6)** | **63** | 51.8 (VIII) | **68** | 0.28 (0.13) |
|  | *SPO13-THR1* (VIII) | 462 | 52 | 0 | 514 | **5.1 (0.7)** | **67** |  |  | ND |
|  | *THR1-LYS2* (VIII) | 292 | 224 | 1 | 517 | **22.2 (1.2)** | **75** |  |  | 0.06 (0.06) |
| *zip1*[F4A,F5A]<br>(K1309) | *HIS4-CEN3* (III) | 297 | 114 | 3 | 414 | **15.9 (1.6)** | **69** | 72.5 (III) | **68** | 0.61 (0.36) |
|  | *CEN3-MAT* (III) | 300 | 113 | 2 | 415 | **15.1 (1.5)** | **75** |  |  | 0.42 (0.30) |
|  | *MAT-RAD18* (III) | 259 | 148 | 6 | 413 | **22.3 (2.0)** | **61** |  |  | 0.67 (0.28) |
|  | *RAD18-HMR* (III) | 272 | 142 | 3 | 417 | **19.2 (1.7)** | **84** |  |  | 0.37 (0.22) |
|  | *SPO11-SPO13* (VIII) | 326 | 86 | 1 | 413 | **11.1 (1.2)** | **28** | 29.2 (VIII) | **38** | 0.38 (0.38) |
|  | *SPO13-THR1* (VIII) | 373 | 34 | 0 | 407 | **4.2 (0.7)** | **55** |  |  | ND |
|  | *THR1-LYS2* (VIII) | 303 | 101 | 2 | 406 | **13.9 (1.5)** | **47** |  |  | 0.52 (0.37) |
| *zip1*[I18A, F19A]<br>(K1282) | *HIS4-CEN3* (III) | 154 | 52 | 0 | 206 | **12.6 (1.5)** | **47** | 79.8 (III) | **75** | ND |
|  | *CEN3-MAT* (III) | 151 | 61 | 0 | 212 | **14.4 (1.6)** | **72** |  |  | ND |
|  | *MAT-RAD18* (III) | 119 | 83 | 8 | 210 | **31.2 (4.1)** | **86** |  |  | 1.37 (0.52) |
|  | *RAD18-HMR* (III) | 137 | 75 | 3 | 215 | **21.6 (2.8)** | **96** |  |  | 0.68 (0.40) |
|  | *SPO11-SPO13* (VIII) | 147 | 60 | 1 | 208 | **15.9 (2.1)** | **41** | 49.1 (VIII) | **64** | 0.37 (0.37) |
|  | *SPO13-THR1* (VIII) | 171 | 29 | 1 | 201 | **8.7 (1.9)** | **114** |  |  | 1.72 (1.74) |
|  | *THR1-LYS2* (VIII) | 107 | 92 | 1 | 200 | **24.5 (2.2)** | **83** |  |  | 0.12 (0.12) |
| *zip3Δ*<br>(K926) | *HIS4-CEN3* (III) | 322 | 161 | 10 | 493 | **22.4 (2.1)** | **84** | 86.5 (III) | **82** | 1.16 (0.38) |
|  | *CEN3-MAT* (III) | 327 | 167 | 5 | 499 | **19.7 (1.6)** | **87** |  |  | 0.54 (0.25) |
|  | *MAT-RAD18* (III) | 318 | 153 | 9 | 480 | **21.6 (2.1)** | **60** |  |  | 1.13 (0.39) |
|  | *RAD18-HMR* (III) | 306 | 174 | 8 | 488 | **22.8 (2.0)** | **100** |  |  | 0.76 (0.28) |
|  | *SPO11-SPO13* (VIII) | 335 | 119 | 7 | 461 | **17.5 (1.9)** | **45** | 35.7 (VIII) | **47** | 1.49 (0.58) |
|  | *SPO13-THR1* (VIII) | 427 | 44 | 1 | 472 | **5.3 (0.9)** | **70** |  |  | 1.83 (1.83) |
|  | *THR1-LYS2* (VIII) | 369 | 97 | 4 | 470 | **12.9 (1.5)** | **44** |  |  | 1.37 (0.69) |
| *zip3Δ msh4Δ*<br>(AM3658/AM3659) | *HIS4-CEN3* (III) | 351 | 195 | 10 | 556 | **22.9 (1.9)** | **86** | 93.1 (III) | **88** | 0.87 (0.28) |
|  | *CEN3-MAT* (III) | 367 | 194 | 8 | 569 | **21.3 (1.7)** | **106** |  |  | 0.73 (0.26) |
|  | *MAT-RAD18* (III) | 344 | 177 | 12 | 533 | **23.4 (2.1)** | **64** |  |  | 1.24 (0.37) |
|  | *RAD18-HMR* (III) | 330 | 210 | 12 | 552 | **25.5 (2.0)** | **111** |  |  | 0.86 (0.26) |
|  | *SPO11-SPO13* (VIII) | 361 | 179 | 8 | 548 | **20.7 (1.8)** | **53** | 42.4 (VIII) | **56** | 0.83 (0.30) |
|  | *SPO13-THR1* (VIII) | 464 | 64 | 1 | 529 | **6.6 (0.9)** | **87** |  |  | 0.95 (0.95) |
|  | *THR1-LYS2* (VIII) | 392 | 129 | 5 | 526 | **15.1 (1.5)** | **51** |  |  | 1.04 (0.47) |
| *zip3Δ zip1*[Δ2-9]<br>(MP52) | *HIS4-CEN3* (III) | 354 | 155 | 5 | 514 | **18.0 (1.6)** | **67** | 80.6 (III) | **76** | 0.78 (0.24) |
|  | *CEN3-MAT* (III) | 355 | 171 | 7 | 533 | **20.0 (1.7)** | **100** |  |  | 0.78 (0.30) |
|  | *MAT-RAD18* (III) | 345 | 159 | 6 | 510 | **19.1 (1.7)** | **53** |  |  | 0.75 (0.31) |
|  | *RAD18-HMR* (III) | 317 | 187 | 9 | 513 | **23.5 (1.9)** | **103** |  |  | 0.77 (0.27) |
|  | *SPO11-SPO13* (VIII) | 343 | 150 | 6 | 499 | **18.6 (1.7)** | **47** | 39.1 (VIII) | **51** | 0.83 (0.35) |
|  | *SPO13-THR1* (VIII) | 438 | 39 | 2 | 479 | **5.3 (1.1)** | **70** |  |  | 4.76 (3.34) |
|  | *THR1-LYS2* (VIII) | 354 | 122 | 4 | 480 | **15.2 (1.6)** | **52** |  |  | 0.85 (0.43) |

**S3 Table.**
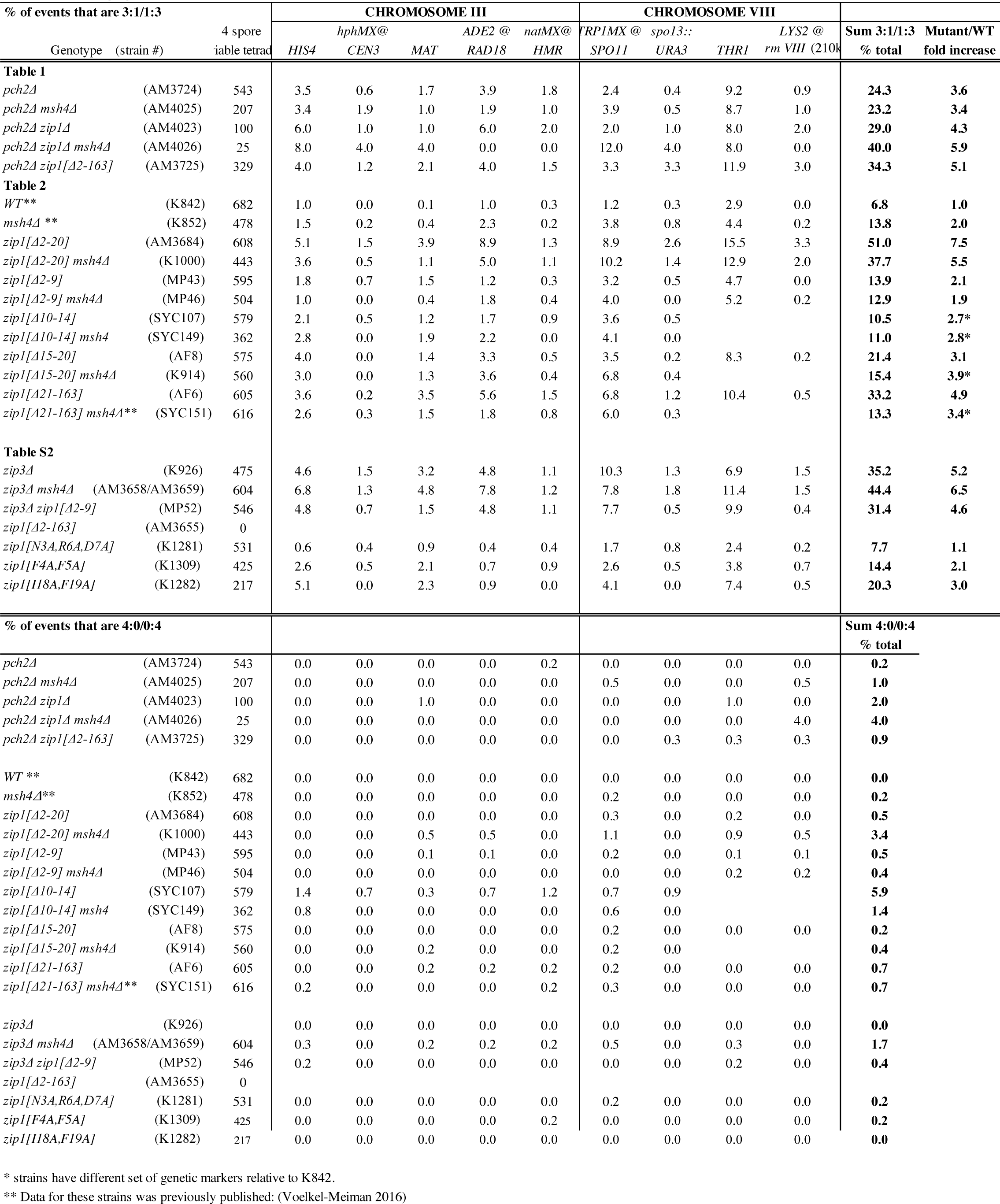
Non-Mendelian segregation (gene conversion events) per locus, measured in 4-spore viable tetrads. Shown are the percentages of non-Mendelian segregation events (3:1/1:3 segregation, top; 4:0/0:4 segregation, below) out of the total tetrads analyzed (second column) in each of the indicated strains. Data is derived from 4-spore viable tetrads with no more than 2 gene conversion (non-2:2) events, although cases where adjacent loci segregate non-2:2 were considered a single conversion event. The sum total percentage of observed non-Mendelian events, and the fold increase relative to wild type, is presented, far right. Strains marked with a single asterisk have a different set of genetic markers on chromosome VIII, relative to the wild-type strain used in this analysis. Strains marked with a double asterisk were previously published[22].

**S4 Table.**
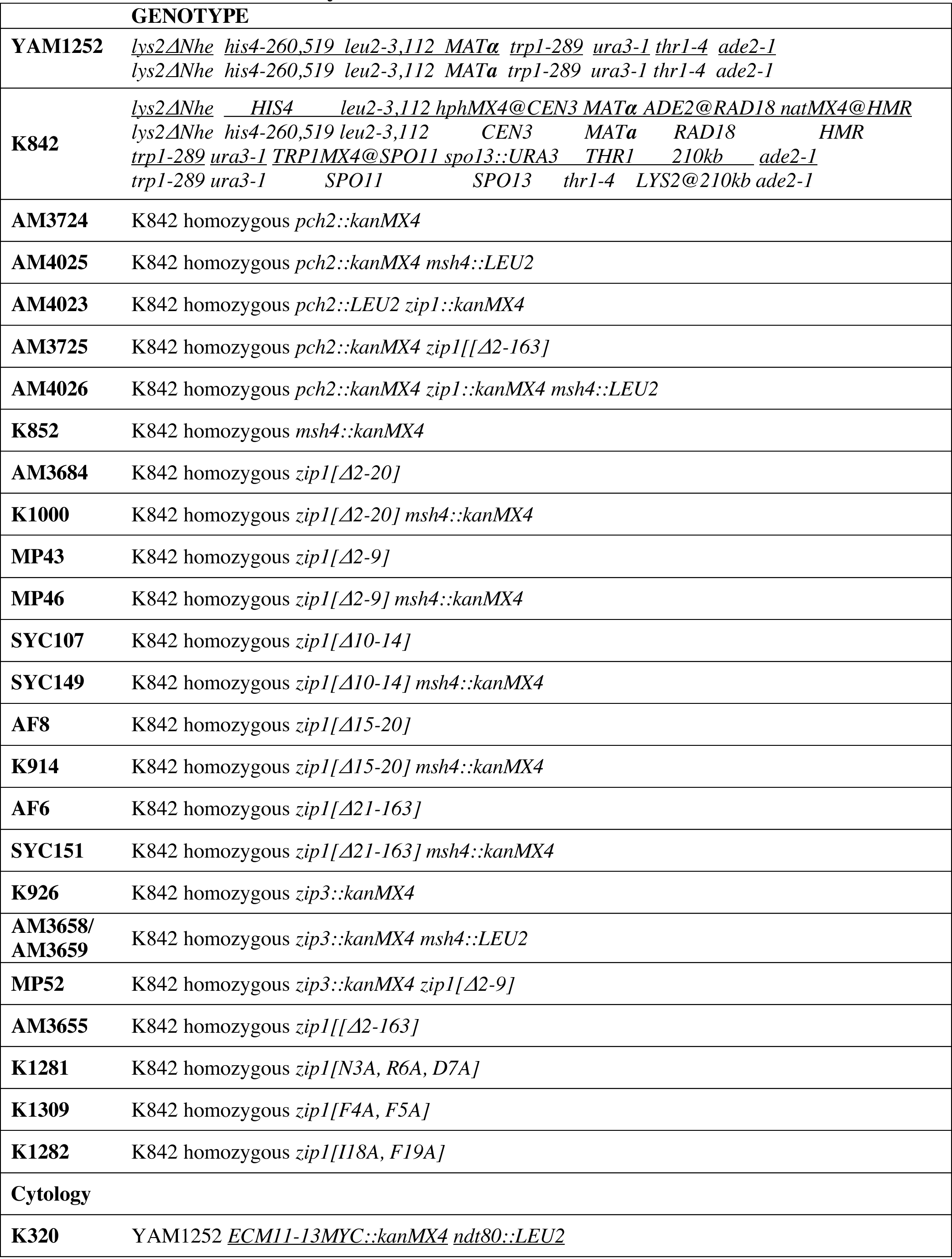
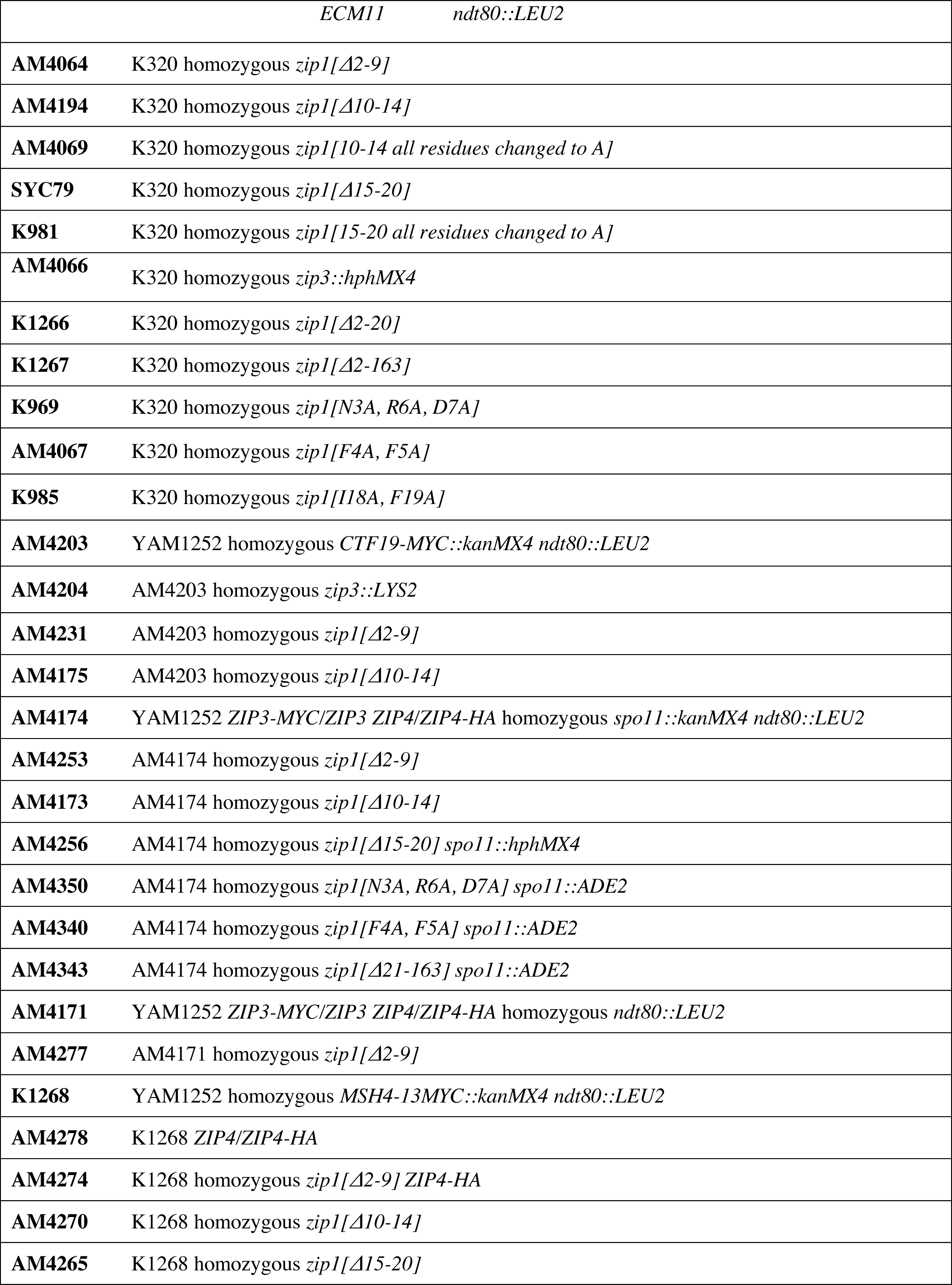
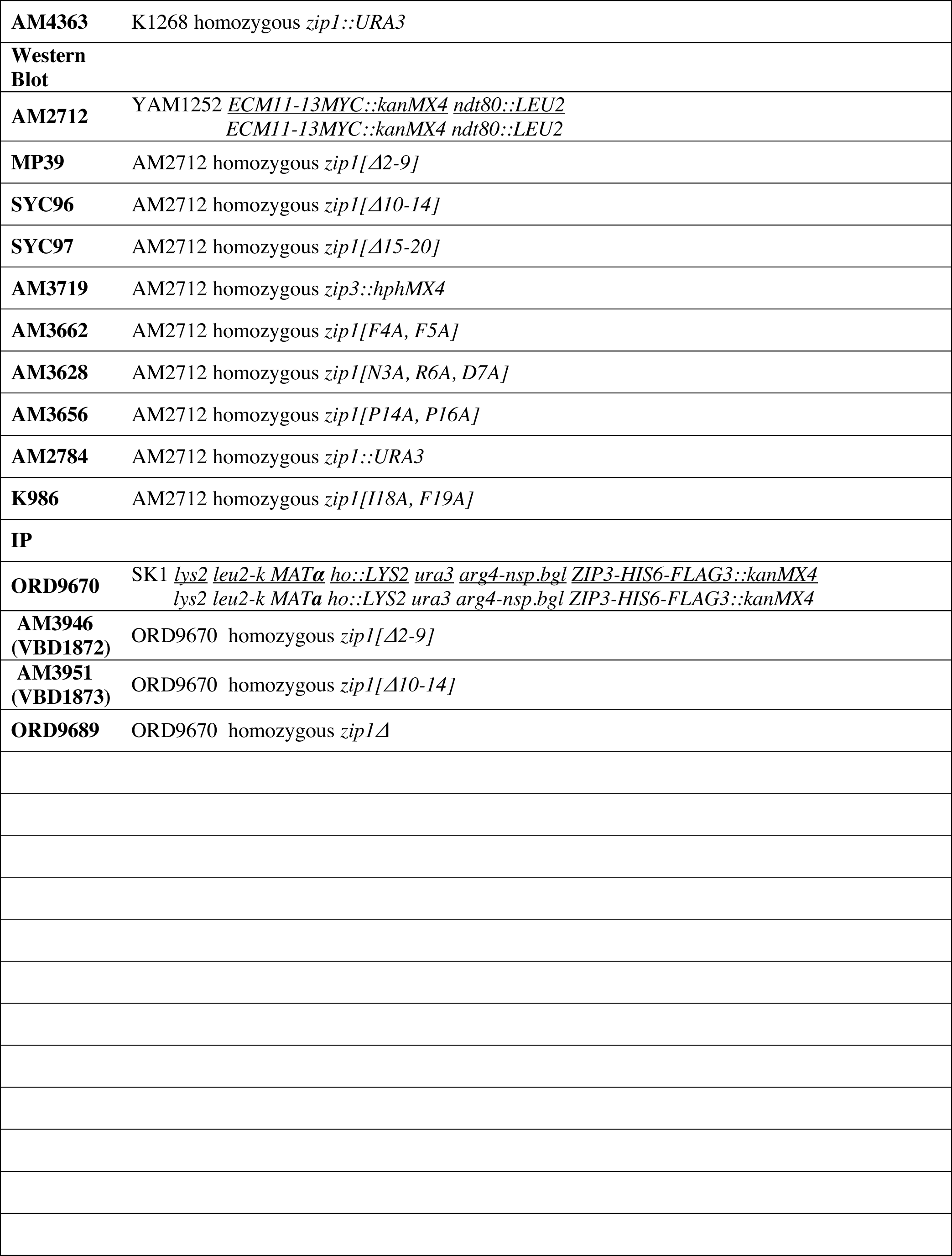
Strains used in this study. Strains are of the BR1919-8B background [52].

## Acknowledgements

We are always grateful for to Dr. Alison North (Director of The Rockefeller University’s Bioimaging Resource Center) for her time and expertise. This work was supported by National Institutes of Health grants 1R15GM104827, and 1R15GM116109 (to A.J.M.). A.dM and V.B. were funded by Institut Curie, CNRS, Labex DEEP (ANR-11-LBX-0044), projet Fondation ARC and La Ligue contre le Cancer. O.R.D. is a Sir Henry Dale Fellow jointly funded by the Wellcome Trust and Royal Society (Grant Number 104158/Z/14/Z) and is also supported by a Royal Society Research Grant (Grant Number RG170118).

